# Meiotic kinetochore reconstitution identifies chromosome segregation factors

**DOI:** 10.64898/2026.09.11.750869

**Authors:** Gerard H. Pieper, Niamh O’Loughlin, Mansour Aboelenain, Lucy Munro, Kousik Sundararajan, Elizabeth Torres-Arce, Darina Lisichkina, Christos Spanos, Tony Ly, Richard A. Anderson, Aaron F. Straight, Adele L. Marston

## Abstract

Kinetochores are complex protein assemblies at centromeres that direct and regulate chromosome segregation^1^. During gamete formation by meiosis, kinetochore specialisations drive two sequential and distinct segregation events to partition half the genome to the offspring^2,3^. Age-dependent segregation errors in human oocytes are a major cause of reproductive failure and are linked to kinetochore dysfunction^4–6^, however the molecular composition and function of mammalian meiotic centromeric chromatin and kinetochores remains poorly defined. Here, we reconstitute vertebrate meiosis I and II centromeric chromatin using meiosis I and II endogenous proteins to uncover conserved regulators of chromosome segregation in mammalian meiosis. We find that, in addition to canonical kinetochore proteins, a specific subset of chromatin regulators is enriched on meiotic centromeric chromatin. We discover previously unrecognized kinetochore proteins conserved from frogs to mouse and humans and demonstrate their localisation at mammalian oocyte centromeres. Among these, we show that the newly identified centromeric protein HSPBAP1, is required to maintain chromosome integrity and ensure faithful segregation in both mouse and human oocytes. These findings define the composition of vertebrate meiotic kinetochores and uncover conserved factors that safeguard mammalian meiotic chromosome segregation, a highly error-prone process and major cause of aneuploidy and reproductive failure in humans^7^.

## Main text

Oocytes preserve and transmit the maternal genome, ensuring its faithful inheritance by the next generation. This occurs through meiosis, a specialized two-step cell division in which chromosomes undergo reductional segregation in meiosis I followed by equational segregation in meiosis II^8^. Chromosome segregation is directed by kinetochores, multiprotein assemblies built on centromeric chromatin that couple chromosomes to spindle microtubules^1^. To execute meiosis, kinetochores must acquire functions distinct from those operating in mitotic cells^2^. Failures in meiotic chromosome segregation are remarkably common in humans and underlie infertility, miscarriage, and clinical syndromes with major effects on health, particularly with increased maternal age when kinetochore function deteriorates through unknown mechanisms^4–7,9,10^.

Oocyte meiosis presents unique challenges for centromere maintenance and kinetochore function. In mammals, oocytes can remain arrested for prolonged periods while preserving centromere identity and chromosome integrity, despite extensive chromatin remodelling during oogenesis^11,12^. Following meiotic resumption, centromeric chromatin must support assembly of kinetochores capable of directing accurate reductional chromosome segregation and ultimately centromere inheritance in the embryo. Maternal centromeric components are also thought to contribute to paternal centromere function after fertilization^13^. Thus, the composition and function of meiotic kinetochores and the underlying centromeric chromatin are of central importance, yet remain poorly understood.

To biochemically define vertebrate meiosis I and II kinetochores, we adapted an *in vitro* assembly system based on synthetic chromatin templates and *Xenopus laevis* egg extracts^14,15^. Chromatin templates comprised either centromere-specific histone variant CENP-A-containing nucleosomes, which templates kinetochore assembly^14,15^, or canonical H3-containing nucleosomes, which templates non-centromeric chromatin^16^. To generate these templates, purified *X. laevis* histones H2A, H2B, H3 and H4 and *Homo sapiens* CENP-A (Extended Data Fig. 1a) were assembled into centromeric (with CENP-A) and non-centromeric (with H3) nucleosomes, loaded onto a DNA array of 18 times ‘601’ Widom nucleosome placement sites (Extended Data Fig. 1b)^17^ and bound to magnetic beads. We first assembled kinetochores in the well-established Xenopus egg extract system, naturally arrested in meiosis II (Fig. 1a) The conserved meiosis-specific kinetochore factor MEIKIN^3,18^, previously uncharacterized in Xenopus, together with the core kinetochore protein CENP-C^19^, were enriched on CENP-A chromatin relative to H3 chromatin, confirming assembly of meiotic kinetochores (Extended Data Fig. 2a, b). To broadly define the protein composition of meiosis II kinetochores, we analysed the eluates from meiosis II extract-incubated CENP-A, H3 and control “no chromatin” beads by mass spectrometry. Pairwise comparisons allowed proteins to be classified as enriched on CENP-A or H3 chromatin or shared (Fig. 1b). As expected, core kinetochore proteins were primarily enriched on CENP-A chromatin (Fig. 1c, d). Almost all kinetochore subcomplexes were represented (Fig. 1c, Extended Data Fig. 2c), including MEIKIN, suggesting the presence of complete kinetochores, and most were specifically detected on CENP-A chromatin (Extended Data Fig. 3). In addition to known centromere-associated proteins such as the FACT complex^20^ (SSRP1 and SUPT16H), novel candidate centromeric proteins including kinase CDK5, ubiquitin ligase UBR4 and JmjC domain protein HSPBAP1 were enriched on CENP-A chromatin. Proteins enriched on both CENP-A and H3 chromatin (“shared”, Fig. 1b) were strongly enriched for DNA repair, DNA replication and chromosome organisation functions (Fig. 1d; Extended Data Fig. 2d), suggesting that their recruitment may be driven by common features of the synthetic chromatin, including the presence of unprotected DNA ends. Both types of chromatin also showed enrichment of condensin I, while condensin II was enriched only on CENP-A chromatin, as expected^21^ (Extended Data Fig. 2e). Chromatin remodellers were also found on both chromatin types, with some showing specificity for CENP-A or H3 chromatin, including the expected H3 nucleosome-enriched HELLS and CDCA7^16^ (Fig. 1c, d). Therefore, in vitro assembly of meiosis II kinetochores on reconstituted centromeric chromatin reveals known and novel centromere-associated proteins.

**Figure 1.**
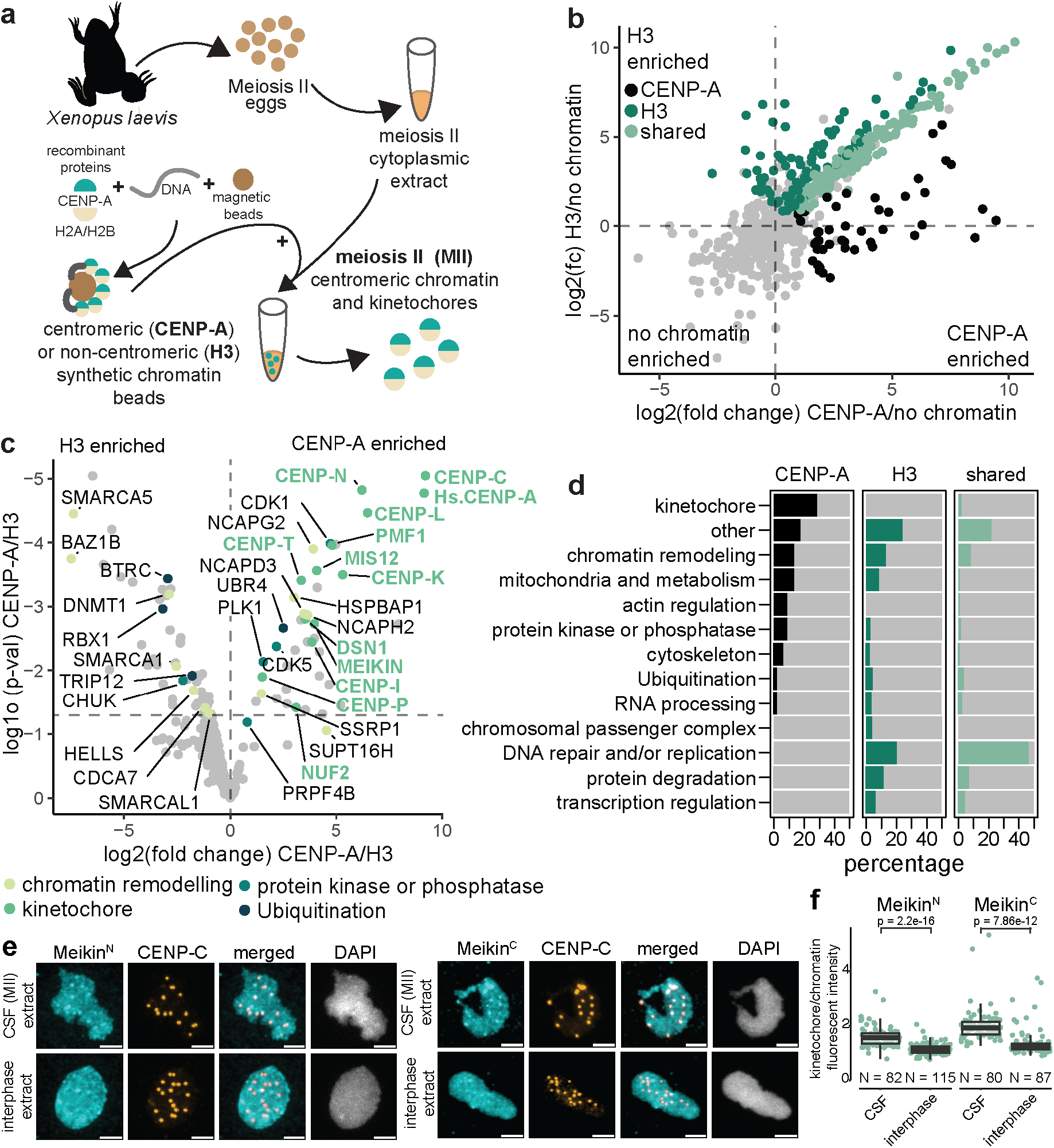
Reconstituted meiosis II centromeric chromatin proteome identifies novel CENP-A associated proteins. **A)** Schematic overview of meiosis II centromeric chromatin reconstitution **B)** Classification of proteins on meiosis II chromatin based on enrichment of CENP-A/no chromatin (X-axis) and H3/no chromatin (Y-axis). **C)** Volcano plot of CENP-A/H3 enrichment (X-axis) and p-values (Y-axis). Labelled proteins in CENP-A and H3 clusters based on protein function. **D)** Annotated biological function representation in meiosis II CENP-A, H3 and shared chromatin proteins. **E)** Chromatin reconstitution in meiosis II (CSF) extract or interphase (CSF+ CaCl2) extract using frog sperm DNA as a template. Nuclei were fixed and immunostained with indicated antibodies. Scale bar represents 5uM. **F)** Quantification of MEIKIN^N^ and MEIKIN^C^ fluorescent intensity on kinetochores over chromatin signal on sperm chromatin assembled in CSF or interphase extract.

MEIKIN was notable among the proteins enriched on MI and MII centromeric chromatin. Mouse MEIKIN is a meiosis-specific regulator of PLK1, which like other members of the MOKIR (<u>M</u>eiosis <u>O</u>ne <u>KI</u>netochore <u>R</u>egulator) family^22^ directs the specialised meiotic segregation pattern^3^. Although MEIKIN was originally thought to function exclusively in meiosis I³, a C-terminal fragment of exogenous MEIKIN persists at mouse oocyte meiosis II kinetochores following its separase-dependent cleavage in anaphase I^18^. Our finding that endogenous MEIKIN associates with reconstituted centromeric chromatin in both MI and MII frog extracts further supported the idea that MEIKIN is not restricted to meiosis I, but raised the question of whether the MII protein is full-length or cleaved. To test this, we raised antibodies to predicted N and C-terminal fragments of frog MEIKIN, separated by a predicted separase cleavage site (Extended Data Fig. 4a) and confirmed their specificity (Extended Data Fig. 4b). Using sperm DNA as a template for in vitro kinetochore assembly in MII extract, we found that both anti-MEIKIN-C and anti-MEIKIN-N sera recognised endogenous MEIKIN on CENP-C-marked kinetochores (Fig. 1e). When MII extract was driven into interphase by the addition of CaCl_2_, MEIKIN could no longer be detected on kinetochores (Fig. 1e, f), indicating that MEIKIN kinetochore localisation is restricted to M-phase. Thus, either full-length MEIKIN or both potential MEIKIN cleavage products can be incorporated into in vitro assembled meiosis I and II kinetochores. Given that the kinetochore-targeting domain resides within the C-terminal region of MEIKIN^3,18^, detection of the N-terminal region at meiosis II kinetochores strongly supports the presence of full-length protein.

To further test whether full-length MEIKIN persists into MII, we injected prophase oocytes with mRNA encoding for N-terminally tagged 6Myc-MEIKIN and followed MEIKIN by anti-Myc western blotting throughout the MI-MII transition (Extended Data Fig. 4d). Interestingly, although a full-length 6Myc-MEIKIN persisted until the end of the time-course, a lower abundance ∼40kDa species, which roughly corresponds to the size of the predicted 6Myc-MEIKIN-N fragment, appeared at the same time as Cyclin B2 levels started to drop at anaphase I (Fig. 2a; Extended Data Fig. 4d). Indeed, this band was not observed upon injection of mRNA encoding a version of 6Myc-MEIKIN carrying two point-mutations to disrupt the putative separase recognition motif (Extended Data Fig. 4a, e). This suggests that *X. laevis* MEIKIN can be cleaved by separase, however only a minor fraction appears to be cleaved. We next asked if N and C-terminal MEIKIN potential fragments remain stable into MII, or are degraded, since the N-terminal fragment contains a degron motif (Extended Data Fig. 4a). Expression of MEIKIN N and C-term fragments alone revealed that the C-terminal fragment is post-translationally modified during meiosis I, like in mouse^23^, however, neither fragment was degraded during the MI-MII transition (Extended Data Fig. 4f, g). Taken together, our findings suggest that although a minor fraction is cleaved, full-length MEIKIN is the predominant form present at meiosis II kinetochores.

**Figure 2.**
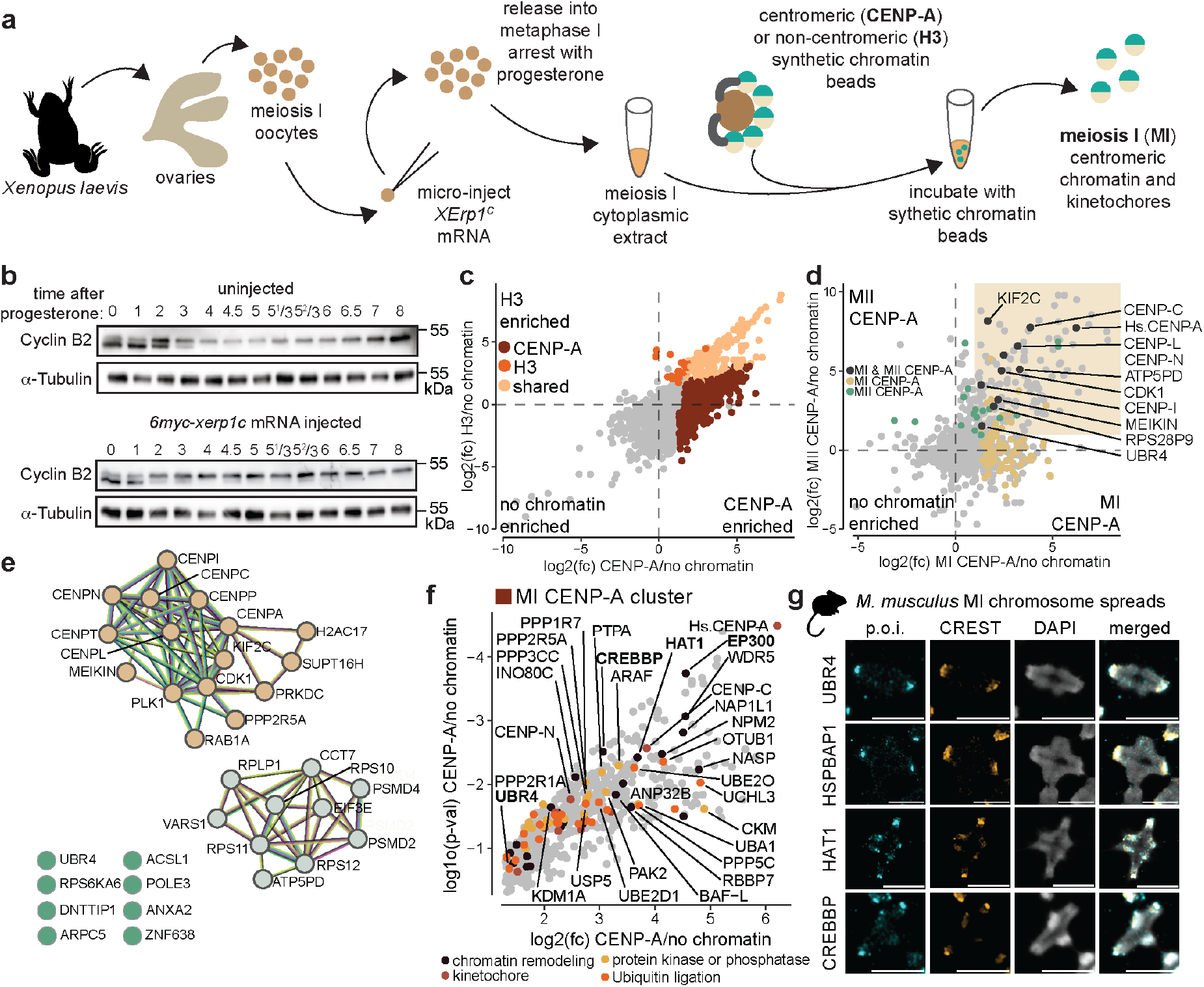
Identification of novel conserved centromeric proteins. **A)** Schematic overview of meiosis I centromeric chromatin reconstitution. **B)** Example of western blot timecourse of 6myc-xerp1c mRNA injected *X. laevis* oocytes and uninjected controls. **C)** Classification of proteins on meiosis I chromatin based on enrichment of CENP-A/no chromatin (X-axis) and H3/no chromatin (Y-axis). **D)** Classification of proteins that are present on both meiosis I and II reconstituted chromatin. Meiosis I CENP-A/no chromatin enrichment on X-axis and meiosis II CENP-A/no chromatin enrichment on the Y-axis. Yellow box represents proteins with a fold change of 2 or higher on both meiosis I and II CENP-A chromatin. **E)** STRING analysis of the proteins enriched on either MI CENP-A, MII CENP-A or both MI and MII CENP-A chromatin highlighted in the yellow box in (D). Natural clusters were found by MCL-clustering with inflation parameter of 3. Brown: protein cluster associated with amplification of signal from unattached kinetochores via MAD2 inhibitory signal. Grey: protein cluster associated with formation of 40S ribosomes. Blue: unclustered. **F)** All proteins in the meiosis I CENP-A cluster, colour coded for protein function. Top 30 most enriched proteins labelled. **G)** Representative mouse oocyte meiosis I chromosome spreads immunostained with indicated antibodies. Scale bar represents 5μm.

We next sought to reconstitute meiosis I kinetochores. Frog oocytes do not arrest in meiosis I and obtaining pure populations by staging oocytes is challenging due to poor synchrony^24^. To overcome this, we set up a robust meiosis I oocyte extract system by induction of metaphase I arrest (Fig. 2a). We used a truncated, constitutively active version of the APC inhibitor XErp1/Emi2^25^, XErp1C, which also harbours two phospho-null mutations (T545A and T551A), rendering it insensitive to inhibition by Cdk1^26,27^. Microinjection of mRNA encoding *XErp1C* into prophase-arrested oocytes resulted in XErp1C production but did not affect kinetics of progesterone-dependent release (Extended Data Fig. 5a, b). Uninjected oocytes progressed through the meiosis I-II transition as observed by the degradation and resynthesis of Cyclin B2^24,28,29^ (Fig. 2b), while Cyclin B2 was stabilised in XErp1C-producing oocytes, indicating successful meiosis I arrest (Fig. 2b). In further experiments, meiosis I-arrested extract was prepared from XErp1C-injected oocytes once 80% had undergone germinal vesicle breakdown (GVBD). Both MEIKIN and CENP-C were enriched on CENP-A chromatin reconstituted in meiosis I extract by immunofluorescence (Extended Data Fig. 5c, d), confirming kinetochore assembly.

To determine the meiosis I kinetochore proteome, we analysed proteins eluted from meiosis I chromatin beads by mass spectrometry (Fig. 2a, c). Proteins were grouped based on enrichment on CENP-A, H3, or both chromatin bead types (Fig. 2c). Multiple core and accessory kinetochore proteins were enriched on CENP-A chromatin (Extended Data Fig. 5e). Among proteins shared between chromatin types, DNA damage proteins and condensin subunits showed the strongest enrichment (Extended Data Fig. 5f), consistent with findings from chromatin assembled in meiosis II extract. However, chromatin assembled in MI extract was associated with substantially more proteins than chromatin assembled in MII extract (Fig. 1b, 2c, Extended Data Fig. 5g), possibly reflecting differences resulting from the two preparation methods. We identified 171 proteins common to both MI and MII CENP-A chromatin (Extended Data Fig. 5g) and found them to be enriched with proteins with chromatin functions, while MI unique proteins were associated with a broader range of functions (Extended Data Fig. 5h). Therefore, despite potential non-specific interactors arising from the lower purity of MI extracts, MI- and MII-assembled chromatin arrays assemble similar proteins, supporting the robustness of the system.

Eleven proteins were significantly enriched on CENP-A chromatin compared to H3 chromatin in both MI and MII (Fig. 2d). These were primarily kinetochore-associated proteins including meiosis-specific MEIKIN (Fig. 2d). Expanding the analysis to all proteins enriched (log2 fold change of at least 1) on CENP-A chromatin in both MI and MII revealed two major clusters (Fig. 2e). The largest cluster represented core kinetochore and kinetochore-associated proteins such as FACT subunits and regulatory kinases and phosphatases including CDK1, PLK1 and PP2A (Fig. 2e). Multiple proteins, including ubiquitin ligase UBR4, could not be sorted into any cluster, suggesting novel kinetochore association of these proteins. Therefore, together, the development of MI-arrest extracts and the robust reconstitution of kinetochores in both MI and MII provide a versatile approach to identify centromeric chromatin-associated proteins across meiotic stages.

Among proteins enriched on MI CENP-A chromatin were enzymes involved in chromatin remodelling, phosphorylation, dephosphorylation or ubiquitination (Fig. 2f). Many of these proteins are implicated in centromere, kinetochore or CENP-A-related function, such as EP300/P300/KAT3B^30^, HAT1/KAT1^31,32^, RBBP7^33^ and NASP^34,35^, again highlighting the specificity of reconstituted centromeric chromatin. To determine whether proteins associated with MI or MII frog centromeric chromatin could have conserved functions at meiotic kinetochores, we first examined the localisation of a subset on mouse oocyte chromosome spreads (Fig. 2g, Extended Data Fig. 6). This confirmed the conserved centromeric enrichment of E3 ligase UBR4^36^, JmjC domain protein HSPBAP1^37^, lysine acetyl transferases HAT1/KAT1 and CREBBP/KAT3A^38^, while lysine acetyl transferase P300/KAT3B^30,38^ was spread throughout the chromatin in both in meiosis I and II (Fig. 2g, Extended Data Fig 6). Overall, reconstitution of Xenopus MI and MII kinetochores identified diverse meiotic kinetochore-associated proteins, including previously unrecognised factors with conserved localisation at mammalian oocyte kinetochores.

As proof of principle that novel proteins identified on reconstituted kinetochores include functional regulators of chromosome segregation, we focused on two novel kinetochore proteins. Firstly, given the established roles of ubiquitin signalling in centromere maintenance, cell-cycle control, and meiotic chromosome organisation^39–41^, we focused on the atypical E3 ubiquitin ligase UBR4. We confirmed that UBR4 was localised to chromosomes with enrichment at kinetochores also in human MI and MII oocytes (Fig. 3a). The levels of human UBR4 enrichment at kinetochores over chromosomes was variable (Fig. 3b), with no UBR4 kinetochore enrichment in several oocytes (kinetochore:chromosome ratio ∼1), but strong enrichment in others. We hypothesised that UBR4 kinetochore enrichment may be linked to the extent of chromosome alignment on the metaphase plate. Indeed, oocytes with low UBR4 enrichment exhibited lower levels of chromosome alignment (Fig. 3c, d). The cell cycle-dependent kinetochore localisation of UBR4 suggested a potential role in cell cycle regulation. To test this, we depleted UBR4 by TRIM-away^42,43^ in mouse oocytes. Although we did not see an effect on meiosis I duration (Extended Data Fig. 7a), we observed markedly delayed GVBD, indicating slower entry into meiosis I (Fig. 3e). Thus, UBR4 is a novel meiotic kinetochore protein conserved from frog to mouse and human with a function in cell cycle regulation.

**Figure 3.**
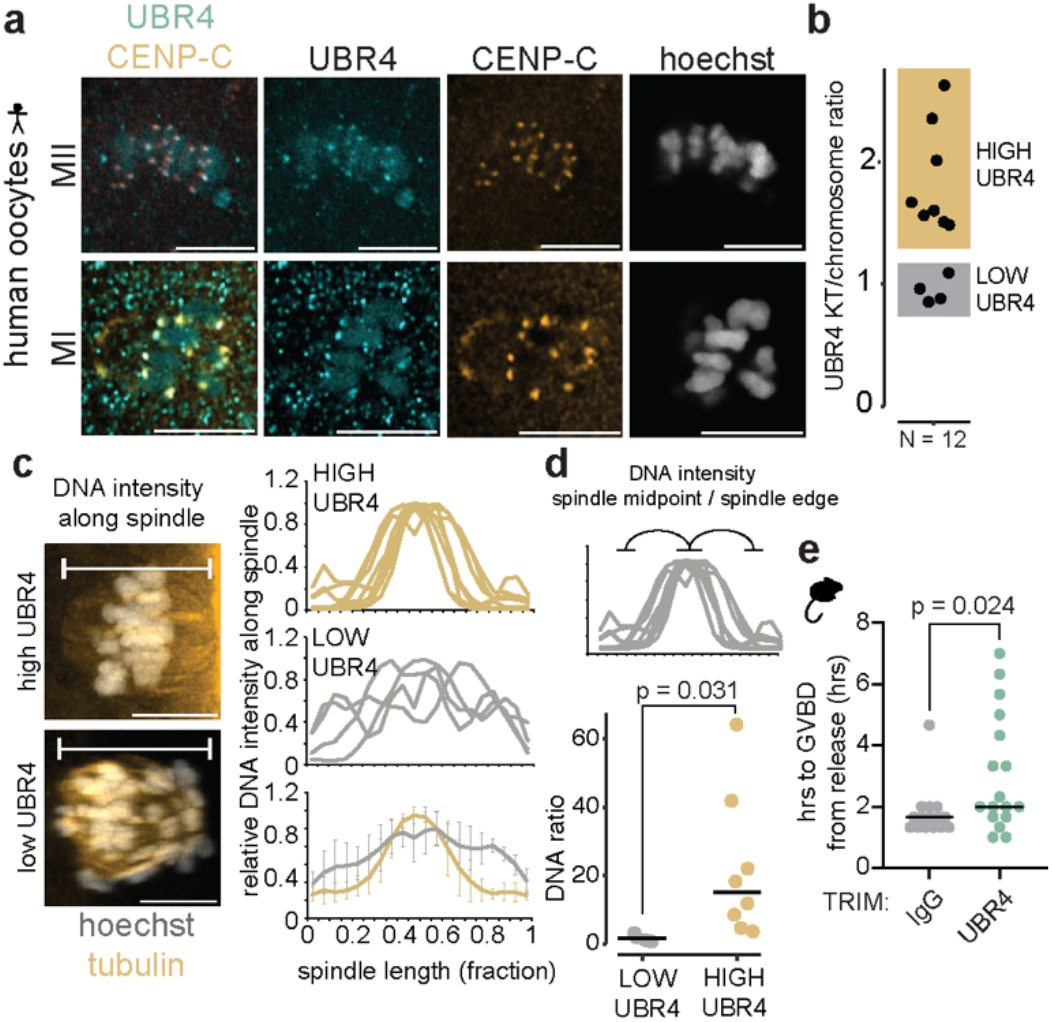
UBR4 is a novel human meiotic kinetochore protein. **A)** Example immunofluorescence images stained with indicated antibodies, showing the different classes of UBR4 kinetochore localisation in human MI (N = 2 from 2 donors) and MII oocytes (N=10 from 7 donors). **B)** Quantification of UBR4 kinetochore over chromosome signal in pooled MI and MII human oocytes shown in (A). **C)** Left: example of 3D representations of human oocyte spindles and chromosomes with either high or low UBR4. Indicated is the DNA measurement area along the spindle. Right: Quantification of normalised DNA signal along the spindle (normalised to spindle length) for high and low UBR4 groups. Bottom graph represents the average of DNA intensity along the spindle for both groups. Error bars represent standard deviation. **D)** Quantification of the ratio of the DNA intensity along the spindle midpoint (bins 0.45-0.55 over the spindle edges (bins 0.1-0.2 and 0.8-0.9) for the UBR4 high (N = 4) and low (N = 8) groups. P-value calculated by t-test. **E)** Timing of GVBD-anaphase I from release of milrinone in mouse oocytes injected with mRNA encoding for *TRIM21-SNAP* and either IgG (N = 17 from 3 replicates) or a-UBR4 antibodies (N = 17 from 3 replicates). P-value calculated by t-test.

Secondly, we focused on HSPBAP1, an uncharacterised JmjC (histone demethylase) domain containing protein^37^ that we found to be enriched on MII CENP-A chromatin. Since HSPBAP1 was not detected by mass-spectrometry on MI chromatin, though its putative interactors KRR1 and RPS29^44^ were (Extended Data Fig. 7b), we confirmed HSBAP1 enrichment on MI CENP-A chromatin beads by immunofluorescence (Extended Data Fig. 7c). Therefore, HSPBAP1 associates with both MI and MII reconstituted *X. laevis* kinetochores, consistent with its presence on both MI and MII mouse oocyte kinetochores (Fig. 2g, Extended Data Fig. 6). To assess HSPBAP1 function, we knocked it down by Trim-away^42,43^ and performed live-imaging of meiosis I chromosome segregation in the presence of SiR-DNA and SPY-tubulin dyes (Fig. 4a). Whereas ∼80% of control oocytes showed normal or near-normal chromosome segregation, we observed pronounced chromosome segregation defects in 50% of HSPBAP1 knockdown oocytes (Fig. 4a, b). In severe cases, a large fraction of chromosomes remained at the metaphase plate after anaphase I, which resulted in chromosome bridges between the polar body and oocyte (Fig. 4c). Even though meiosis I duration between controls and HSPBAP1 knockdown was not significantly different overall (Extended Data Fig. 7d), HSPBAP1 knockdown oocytes which showed abnormal and severe chromosome segregation phenotype were significantly delayed (Fig. 4d). These observations were further recapitulated in human oocytes.

**Fig. 4.**
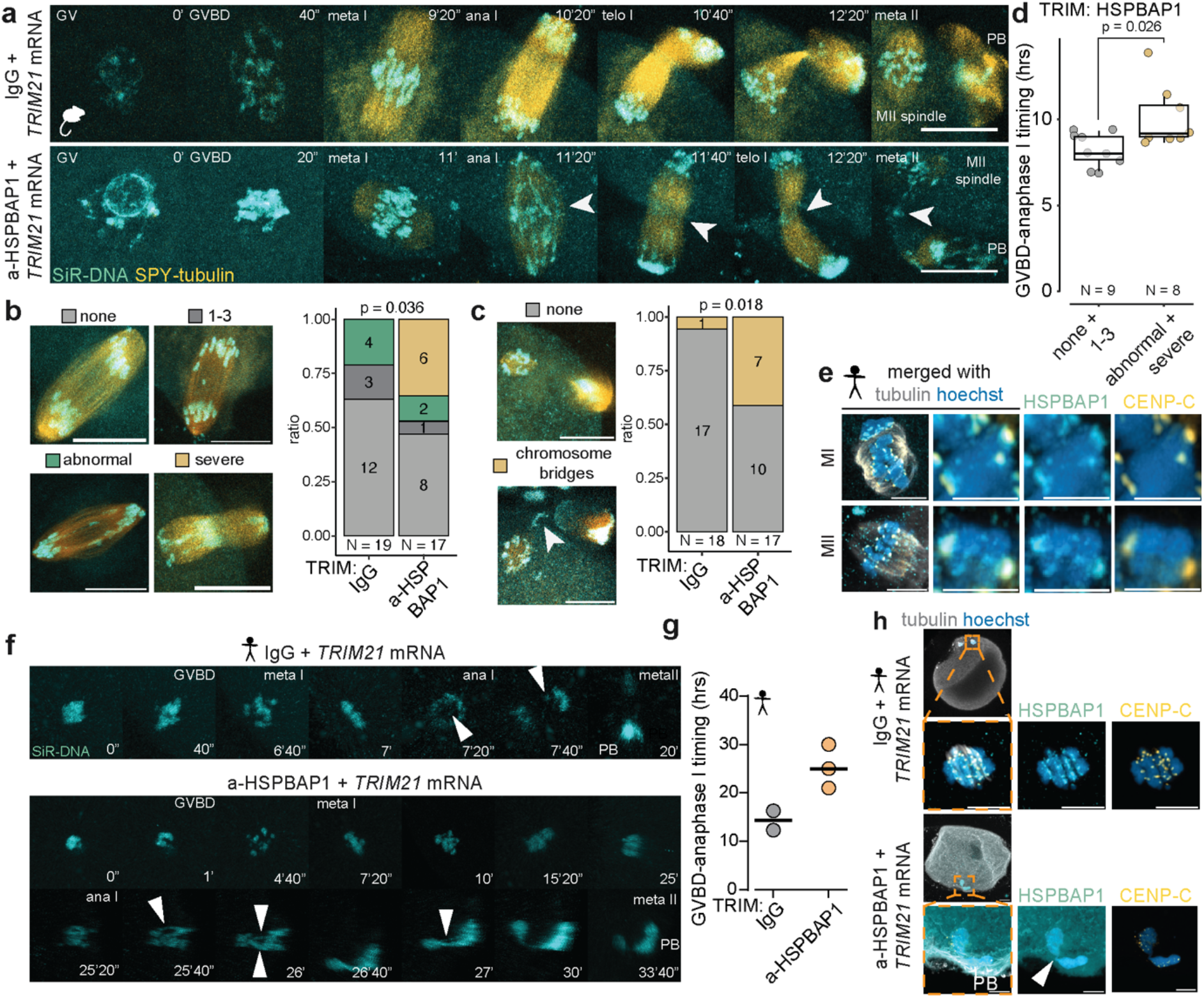
HSPBAP1 is a novel mammalian meiotic centromeric protein that functions in chromosome segregation. **A)** Representative images of microinjected mouse oocytes with *Mm.TRIM21-SNAP* mRNA and either rabbit IgG or HSPBAP1 antibodies followed by live imaging of meiosis with SiR-DNA and SPY-tubulin live-imaging dyes. Arrowheads indicate lagging chromosomes or chromosome bridges. Scale bar represents 20μm. **B)** Quantification of lagging chromosomes in HSPBAP1 TRIM-away mouse oocytes. Scale bar in representative images represents 20μm. P-value calculated by Fisher’s exact test. **C)** Quantification of chromosome bridges in HSPBAP1 TRIM-away mouse oocytes. Scale bar in representative images represents 20μm. P-value calculated by Fisher’s exact test. **D)** Quantification of the GVBD-anaphase I timing of micro-injected mouse oocytes shown in (A). 4 biological repeats with a total of 19 and 17 oocytes imaged for IgG and HSPBAP1 trim-away respectively. P-value calculated by t-test. **E)** Immunostaining of human meiosis I (N=4) and meiosis II (N=5) oocytes with indicated antibodies. Maximum Z-stack of several Z-slices through the centre of the spindle are shown. Scale bar indicates 5μm. Single chromosome of each spindle are shown, scale bar indicates 2μm. **F)** Representative 3D Imaris models of microinjected human oocytes with *Hs.TRIM21-eGFP* mRNA and either rabbit IgG (N=2) or HSPBAP1 (N=3) antibodies followed by live imaging of meiosis with the DNA dye SiR-DNA live-imaging dye. Arrowheads indicate lagging chromosomes or chromosome bridges. 2 oocytes for each group from a single patient. **G)** Quantification of the GVBD-anaphase I timing of micro-injected human oocytes shown in (G). **H)** Immunostaining of fixed HSPBAP1 TRIM away human oocytes shown in (G) with indicated antibodies. Whole maximum Z-projections are shown. Scale bar in whole oocytes represents 10μm, scale bar in zoomed in images represents 5μM.

HSPBAP1 localised to centromeres in both MI and MII (Fig. 4e) and knockdown of HSPBAP1 by TRIM-away replicated the chromosome segregation-defect observed in mouse (3/3 human oocytes severe lagging chromosomes, 2/3 chromosome bridges; Fig. 4f). All human oocytes where HSPBAP1 knockdown was knocked down exhibited a strong delay in meiosis I duration (Fig. 4g), similar to our findings in mouse (Fig. 4d). In two out of three human HSPBAP1 Trim-away oocytes chromosomes were clustered together in both the polar body and the oocytes, maintaining chromosome bridges (Fig. 4h). These two phenotypes suggest that HSPBAP1 might be involved in chromosome condensation and establishing correct microtubule-kinetochore attachments in both mouse and human. Thus, we have identified a novel centromere-localised JmjC domain protein involved in regulating chromosome segregation in meiosis I that is conserved from frog to mammals including humans. Other histone demethylases with functions at the centromere appear to be involved in regulating the maintenance of centromeric heterochromatin and the regulation of centromere transcription^45,46^. However, HSPBAP1, as a CENP-A chromatin/kinetochore associated protein, might have different and unique functions at meiotic centromeres, potentially related to kinetochore function.

In conclusion, here we have developed methods allowing for the reconstitution and proteomic analysis of centromeric chromatin in meiosis. Using native proteins from meiosis I and II *X. laevis* oocytes we provide, for the first time, insight into the composition of meiotic centromeres, overcoming technical barriers that conventional whole cell-based methods bring. This resulted in the identification of several novel meiotic centromere-enriched proteins conserved from frog to mammals. In addition, we found novel functions for E3-ligase UBR4 and JmjC domain protein HSPBAP1 in mouse and human meiotic chromosome segregation. Thus, we have expanded the proteomic landscape of meiotic centromeres to include new classes of proteins. It will be key to investigate whether dysfunction of these proteins is related to age-dependent chromosome segregation defects in human oocytes and if they contribute to mediating chromosome segregation in other cellular contexts.

## Acknowledgements

We gratefully acknowledge the Wellcome Discovery Research Platform for Hidden Cell Biology Light Microscopy Core, the European Xenopus Resource Centre (EXRC) at the University of Portsmouth and the School of Biological Sciences Workshop. We thank Ana Losada, Thomas Mayer and Binyam Mogessie for reagents, Ian Adams for supporting the mouse work as Project Procedure Licence holder and Thomas Mayer for advice on Xenopus oocyte arrest and microinjection. We are grateful to Marston lab members, the Eggs and Embryos group for useful discussion and to Weronika Borek for initial exploratory experiments.

## Author contributions

Conceptualization – GHP and ALM; Methodology – GHP, KS, AFS; Formal analysis – GHP, NOL, MA, LM, ETA, DL, CS, TL; Investigation – GHP, NOL, MA, LM, JS, ETA, DL; Resources GHP, MA, LM, RAA; Writing-original draft – GHP; writing-review and editing; GHP and ALM with input from all authors; visualisation – GHP; supervision-ALM, AFS and GHP; Project administration – GHP and ALM; funding acquisition-GHP, ALM, TL.

## Materials and Methods

### Key resources

Key reagents, antibodies, materials and plasmids used in this study are given in Supplementary Table 1.

### *X. laevis* CSF extract preparation

Collection of eggs and CSF extract experiments were performed at the European Xenopus Resource Centre (EXRC) at the University of Portsmouth. Females were primed with PMSG (pregnant mare serum gonadotrophin) and were injected with 600U HCG (human chorionic gonadotrophin 1500 U/ml) the night before the experiment. Frogs were kept in 100mM NaCl overnight. The following morning, eggs were transferred into glass dishes with MMR (100 mM NaCl, 2 mM KCl, 1 mM MgSO_4_, 2 mM CaCl_2_, 5 mM HEPES, 0.1 mM EDTA, pH 7.8 with NaOH). Eggs were dejellied with fresh 2% cysteine in water (pH 7.8) by gentrly swirling for 5 min, replacing the cysteine several ties. Cysteine was washed out gently with MMR and bad eggs removed. Then, eggs were transferred to petridishes with 1xXB, then washed 3x with 1xXB media and 2x with XB supplemented with protease inhibitors (1μg/ml), 5mM EGTA and 1mM MgCl_2_. Eggs were transferred to an Ultraclear tube with 1ml of 1xXB supplemented with 10μg/ml protease inhibitors and 10μg/ml cytochalasin D. Eggs were packed by centrifugation at 1000RPM for 60 sec and then at 2000RPM for 60 sec and excess buffer was removed. Eggs were crushed by centrifugation at 10.000RPM at 16°C in a SW41 rotor. Cytoplasm was removed using a 1ml syringe and an 18 gauge needle inserted above the pigment/yolk bottom layer. The cytoplasm (CSF extract) was supplemented with protease inhibitors (final concentrations: 10μg/ml leupeptin, 10μg/ml pepstatin-A, 5μg/ml chymostatin), cytochalasin D (final concentration 10μg/ml) and sucrose (final concentration 50mM) and transferred to ice.

### *X. laevis* meiosis I oocyte extract preparation

*X. laevis* ovaries were obtained from the EXRC. Oocytes were extracted firstly by manually tearing the oocytes in 0.5mm^2^ pieces, then followed by incubation in 2mg/ml collagenase B in MMR media for 50-70 min at room temperature in a 50ml tube, rotating end over end. Collaganese was washed out in 10 washes with MMR. Stage VI oocytes were manually sorted. Then, stage VI oocytes were injected with 10ng of *XErp1C-6myc* mRNA Oocytes were moved to recovery media (MBS (80mM NaCl, 1mM KCl, 1mM MgSO_4_.7H20, 5mM HEPES, 25mM NaHCO_3_, 0.7mM CaCl_2_, pH7.8) + 3% Ficoll) and allowed to express protein overnight at 18°C. Next day, oocytes were incubated with 10μg/ml progesterone in MBS + 3% Ficoll at room temperature and monitored every 30 min for the appearance of a white spot on the animal (dark) pole, which marks GVBD. When 80% of oocytes had undergone GVBD (roughly 5.5-6hrs) oocytes were harvested by 3 washes in fresh CSF-XB buffer (1x XB salts, 50mM sucrose, 10mM HEPES pH7.7, 5mM EGTA, 1mM MgCl_2_, for 100ml add 10ul of 10mM NaOH) (20xXB salts: 2M KCl, 20mM MgCl_2_, 2mM CalCl_2_) then packed into 700ul ultra clear tubes with CSF-XB and 0.1μg/ml cytochalasin B by two 5sec spins at 100g, removing the excess buffer. Tubes were placed in adaptors and spun in a SW55-Ti rotor for 20 min at 15.000g at 4°C in a Beckman ultracentrifuge to crush the oocytes. The cytoplasmic layer was harvested by poking a hole in the side of the tube with a needle and extracting the cytoplasm, taking care to take as little of the pigment/yolk (bottom) and lipid (top) layers as possible. If necessary, another spin for 10 min at max speed in a microcentrifuge at 4°C in a prechilled tube was used to clarify the cytoplasmic layer. MI cytoplasmic extract was transferred to ice and energy mix (final concentrations 10mM creatine phosphate, 1mM MgCl_2_, 1mM adenosine triphosphate)^47^, protease inhibitors (final concentrations: 10μg/ml leupeptin, 10μg/ml pepstatin-A, 5μg/ml chymostatin), cytochalasin D (final concentration 10μg/ml) and sucrose (final concentration 50mM) were added to the cytoplasmic extract. Extracts were split into reactions of 20μl and chromatin assembly reactions were performed as for CSF extract (described below).

### Recombinant histone preparation

Performed exactly as described^15^. Recombinant X. laevis histones were expressed in *Escherichia coli* by transforming the relevant plasmid into BL21 bacteria on chloramphenicol/ampicillin plates. Next day, a single colony was inoculated into preculture 5ml of 2xYT (1L:16g tryptone,10g yeast extract, 5g NaCl, Adjust pH to 7.0) media + chloramphenicol/ampicillin and grown overnight at 37°C. The preculture was diluted 1:1000 into 1L of 2xYT +ampicillin and grown at 37°C and 200RPM until an OD_600_ of 0.5-0.6 at which point expression was induced by 0.2mM IPTG. Cultures were grown for a further 3hrs then harvested. Cell pellets were resuspended in 25ml of buffer W (50mM Tris-HCl, 100mM NaCl, 1mM EDTA, 5mM 2-mercaptoethanol, 0.2mM PMSF) and snap frozen in small droplets in liquid N_2_ and stored at −80°C until use.

Cells were then thawed in a 37°C water bath with constant shaking. Then sonicated six times with a probe sonicator 30sec on/1min off at 200W on ice. The lysate was spun down at 25400g for 20 min at 4°C in a JA-20 rotor. The pellet with inclusion bodies was washed 3x with 25ml buffer TW (Add 0.1% (v/v) Triton X-100 to buffer W) and 2x with 25ml buffer W. Cell pellets were thoroughly mixed with 175ul of DMSO, then with 13.2ml of unfolding buffer (7M guanidine - HCl, 20mM Tris-HCl (pH7.5), 10mM DTT, Prepare fresh) and soaked for 1hr at RT with gentle shaking. Pellets were homogenised with a Dounce homogeniser. The homogenate was centrifuged at 17640g for 20 min at 4°C and the supernatant was saved. Pellet was again resuspended in unfolding buffer, centrifuged and the supernatant was combined with the previous supernatant. Supernatant was dialysed overnight into buffer urea buffer. Histones were purified by FPLC using HiTrapQ and S columns, dialysed into H2O with 0.1mM PMSF and 5mM mercaptoethanol, then lyophylised and aliquots were snap frozen in liquid N_2_ and stored at −80°C.

Human CENP-A was coexpressed with *X. laevis* H4 using a bicistronic plasmid in 6Lof 2xYT media as before but expressed for 6hrs and pellet resuspension in PBS before snap freezing. Sonicated pellets were spun down at 110.000g in a Ti70 rotor for 1hr. CENP-A and H4 were purified form the supernatant with an HA column, dialysed into dialysis buffer (10 mM Tris-HCl (pH 7.4), 0.75 M NaCl, 10 mM 2-mercaptoethanol and 0.5 mM EDTA) and purified on a HiTrap SP FF column. Relevant fractions were snap frozen and stored at −80°C.

H3-H4 tetramers and H2A-H2B dimers were assembled by mixing of equimolar ratios and dialysed into refolding buffer. Proteins were injected onto a Superdex 200 16/60 column and fractions containing roughly equal H3/H4 or H2A/H2B were pooled, aliquoted and snap frozen in liquid N_2_.

### Chromatin array preparation

Chromatin arrays were prepared as described previously^15^, except where noted. A plasmid with an array of 18×601, flanked by AvaI restriction sites was digested with EcoRI, XbaI, DraI and HaeII enzymes (All NEB). The 18×601 fragment was purified by serial PEG dilution and dialysed into TE buffer. The plasmid was biotinylated in a reaction with biotin-dATP, biotin-16-dUTP, alpha-thio-dCTP and dGTP (all at 35mM) in 20ul reactions with 2μl of klenow fragment (5U/μl) and 10μg of DNA for 3hrs at 37°C. DNA was column purified and biotinylation efficiency was assessed by incubation with streptavidin-FITC and running the reactions on a 1% agarose gel.

Chromatin arrays were assembled by mixing equimolar or near equimolar concentrations of CENP-A-H4 or H3-H4 with H2A-H2B and 18×601 arrays followed by salt dialysis buffer exchange. Incorporation rate was assessed by AvaI cleavage of chromatin and running on a native acrylamide gel before soaking in 1μl g/ml ethidium bromide. Chromatin arrays were stored at 4°C until use for 3 months.

### *In vitro* chromatin reconstitution

Chromatin was reconstituted as described previously^15^. First chromatin beads were prepared using 1.5μl of beads per 20μl assembly reaction. Beads were first washed in bead buffer, then resuspended in bead buffer + 2.5% PVA at a ratio of 17.5μl bead buffer + PVA/1μl of beads. 1.5μl of chromatin arrays were added to the beads and incubated at RT for 1hr with mild agitation to prevent settling. Beads were washed 2x with CSF-XB and stored on ice until use. CSF-XB buffer was removed from the chromatin beads and 20μl of cytoplasmic extract was added. Reactions were incubated for 75 min at 19°C. Tubes were flicked every 15 minutes to prevent beads from settling. Extracts were diluted with 100μl CSF-XBT (Add Triton to 0.05% (v/v) final concentration) then washed 3x with CSF-XBT, fixed for 5 min in 100μl CSF-XBTF (add formaldehyde to a final concentration 0.2% (v/v) to CSF-XBT), at room temperature. then washed again 3x in CSF-XBT. All buffers except for CSF-XBTF were chilled to 4°C.

For immunofluorescence^48^: Beads were washed 3x in Abdil (150mM NaCl, 20mM Tris-HCl (pH 7.4), 0.1% Triton X100, 2% (w/v) BSA, store at 4dC) then allowed to adhere to poly-lysine-coated coverslips overnight at 4°C in Abdil.

For mass spectrometry: Beads were washed 3x in CSF-XBT, then proteins were eluted from beads by 2 incubations in 25μl 0.1% rapigest in 50mM Tris-HCl pH8.0 for 10 min at 50°C (final eluate 50μl). Samples were snap frozen in liquid N and stored at −80°C until use.

### Immunofluorescence of chromatin beads

Chromatin beads from chromatin assembly reactions were stained for 1hr at RT with primary antibodies 1:100 from stock in Abdil, then washed 3x with Abdil. They were then incubated with secondary antibodies 1:500 for 45 min at RT washed 2x with Abdil and 2x with PBS. Coverslips were then mounted on microscopy slides in vectashield and sealed with nail polish. Slides were kept at −20°C until use.

### Proteomics of reconstituted chromatin

Protein samples from all biological replicates were processed at the same time and with using the same digestion protocol without any deviations. They were subjected for MS analysis under the same conditions. Protein and peptide lists generated using the same software and the same parameters. Specifically, proteins from each sample were digested using the Filter Aided Sample Preparation (FASP) protocol as described^49^ with minor modifications. In brief, each protein sample was added on the top of a 30 kDa MWCO filter units (Vivacon, UK) along with 150 µl of denaturation buffer (8M Urea in 50mM ammonium bicarbonate (ABC) (Sigma Aldrich)) and spun at 14,000 x g for 20 min, while another wash with 200 µl of denaturation buffer was performed under the same conditions. The protein samples were then reduced by the addition of 100 µl of 10 mM dithiothreitol (Sigma Aldrich, UK) in denaturation buffer for 30 min at ambient temperature, and alkylated by adding 100 µl of 55 mM iodoacetamide (Sigma Aldrich, UK) in denaturation buffer for 20 min at ambient temperature in the dark. Two washes with 100 µl of denaturation buffer and two with digestion buffer (50mM ABC) were performed under the same conditions described above before the addition of trypsin (Pierce, UK). The protease:protein ratio was 1:50 and proteins were digested overnight at 37°C. Following digestion, samples were spun at 14,000 x g for 20 min and the flow-through containing digested peptides was collected. Filters were then washed one more time with 100 µl of digestion buffer and the flow-through was collected again. The eluates from the filter units were acidified using 20 µl of 10% Trifluoroacetic Acid (TFA) (Sigma Aldrich), and spun onto StageTips as described^50^. Peptides were eluted in 40 μL of 80% acetonitrile in 0.1% TFA and concentrated down to 1 μL by vacuum centrifugation (Concentrator 5301, Eppendorf, UK). The peptide sample was then prepared for LC-MS/MS analysis by diluting it to 5 μL by 0.1% TFA.

LC-MS analyses were performed on an Orbitrap Exploris™ 480 Mass Spectrometer (Thermo Fisher Scientific, UK) coupled on-line, to an Ultimate 3000 HPLC (Dionex, Thermo Fisher Scientific, UK). Peptides were separated on a 50 cm (2 µm particle size) EASY-Spray column (Thermo Scientific, UK), which was assembled on an EASY-Spray source (Thermo Scientific, UK) and operated constantly at 55°C. Mobile phase A consisted of 0.1% formic acid in LC-MS grade water and mobile phase B consisted of 80% acetonitrile and 0.1% formic acid. Peptides were loaded onto the column at a flow rate of 0.3 μL min^−1^ and eluted at a flow rate of 0.25 μL min^−1^ according to the following gradient: 2 to 40% mobile phase B in 150 min and then to 95% in 11 min. Mobile phase B was retained at 95% for 5 min and returned back to 2% a minute after until the end of the run (190 min).

Survey scans were recorded at 120,000 resolution (scan range 350-1650 m/z) with an ion target of 5.0e6, and injection time of 20ms. MS2 Data Independent Acquisition (DIA) was performed in the orbitrap at 30,000 resolution with a scan range of 200-2000 m/z, maximum injection time of 55ms and AGC target of 3.0E6 ions. We used HCD fragmentation^51^ with stepped collision energy of 25.5, 27 and 30. We used variable isolation windows throughout the scan range ranging from 10.5 to 50.5 m/z. Narrower isolation windows (10.5-18.5 m/z) were applied from 400-800 m/z and then gradually increased to 50.5 m/z until the end of the scan range. The default charge state was set to 3. Data for both survey and MS/MS scans were acquired in profile mode.

The DIA-NN software platform^52^ version 1.9.2. was used to process the raw files and search was conducted against the *Xenopus laevis* database (XenBase - released in January 2025) with the addition of the CENPA human protein sequence (Uniprot). Precursor ion generation was based on the chosen protein database (automatically generated spectral library) with deep-learning based spectra, retention time and IMs prediction. Digestion mode was set to specific with trypsin allowing maximum of two missed cleavages. Carbamidomethylation of cysteine was set as fixed modification. Oxidation of methionine, and acetylation of the N-terminus were set as variable modifications. The parameters for peptide length range, precursor charge range, precursor m/z range and fragment ion m/z range as well as other software parameters were used with their default values. The precursor FDR was set to 1%.

### Analysis of proteomics data

Proteomics data was analysed with and proteomics plots generated with R software (https://www.R-project.org/). MII and MI data sets were analysed separately in the following way: Xenopus proteins from S and L homeologues and different isoforms were summed to create a single value per protein. Zero values were replaced with NAs. Xenopus gene names were transferred to human orthologues by capitalising or manual search for alternative names and homologues based on information provided on Xenbase^53^ and NCBI databases^54^. Data was processed using functions from the DEP^55^ package and associated functions from the limma package^56^. Initial data processing was done with the DEP package. Repeats were analysed by comparing PCA plots, heatmaps and protein number and repeats were rejected based on deviation from other repeats based on these. For MII data, 2 repeats for each bead type were included and for MI data 3 CENP-A and H3 repeats and 2 no chromatin repeats were included. For MII proteins to be included in the MII dataset, they needed to be detected at least on both CENP-A repeats, both H3 repeats or both no chromatin repeats. For MI proteins to be included in the MI data set they needed to be detected at least in all CENP-A repeats, or all H3 repeats or both no chromatin repeats.

Data was normalised by median normalisation. Missing data was imputed using the MinProb function with q = 0.01 and tune.sigma = 1. Proteins eluted from all bead types were compared to all other beads and statistically significant comparisons were generated with alpha = 0.05 and lfc = 1 cutoffs. P-values were log10 transformed. Data was classified as being significantly enriched on CENP-A, H3 or both chromatin types. For CENP-A proteins were included based on three criteria: 1) if they were significant CENP-A/no chromatin and not significant H3/no chromatin. 2) Significant CENP-A/H3 and significant CENP-A/no chromatin. 3) Significant CENP-A/H3 with fold change > 2 and a positive fold change comparing CENP-A/no chromatin >1.3. The last group was included to exclude proteins that have a high significance when comparing CENP-A/H3 but are either low abundance or enriched on no chromatin beads. H3 proteins were classified by similar rules. Shared proteins were included based on three criteria: 1) CENP-A/no chromatin p-value <= −1.3 and H3/no chromatin p-value <= −1.3. 2) A positive fold change comparing CENP-A and H3 chromatin to no chromatin. 3) They were not included in either the CENP-A or H3 groups. Proteins were classified by protein function by manually analysing proteins on genecards.org^57^.

To define in which chromatin bead groups a protein was detected (Ext. Data Fig. 3), we started from the raw protein intensity data. Protein intensity values were averaged between repeats, if a protein was detected in only a single repeat, that value was taken, thus for this analysis, values represent proteins that were detected in at least 1 repeat. Grey squares indicate proteins that were not detected on that type of chromatin bead.

MI and MII chromatin common proteins (Fig. 2d) were analysed by selecting all proteins that were significant on any bead type of either MI or MII chromatin. The Venn diagram of Ext. Data. Fig 2g was generated with the VennDiagram package (DOI: 10.32614/CRAN.package.VennDiagram) and manually coloured with Adobe Illustrator. For further analysis of proteins positive on CENP-A chromatin in both MI and MII proteins were selected if they were classified as CENP-A enriched in either MI or MII and had a fold change of >2. These were then analysed by STRING analysis^58^. One protein (RPS28P9) was excluded due to lack of protein homologue in the human database. MCL clustering was performed with inflation parameter of 3.

The clustered heatmap of Ext. Data Fig. 2e was generated using the pheatmap R package (DOI: 10.32614/CRAN.package.pheatmap) with cutree_rows = 2. Clusters were then retroactively found and labelled by using the hclust and cuttree functions The scale colour palette was adjusted to the log2 transformed fold changes.

To search for kinetochore proteins in the MI data set (Ext. Data Fig. 5e) we screened for proteins from the following GO terms: GO:0000776 kinetochore, GO:0000775 centromeric region, GO:0034501 kinetochore localisation, GO: 0007052 mitotic spindle organisation, GO: 0071173 spindle assembly checkpoint, GO:0000159 protein phosphatase type 2A (PP2A) complex, GO:0072357 PP1 phosphatase complex. Proteins were colour coded for manually annotated functions.

GO term analysis (Ext. Data Fig. 5h) was done on the proteins of MI only, MII only and MI and MII shared proteins in Ext. Data Fig. 5g using gprofiler^59^ with standard multiquery settings.

### *X. laevis* sperm DNA reactions and immunofluorescence

Sperm DNA reactions were performed at Stanford University. Mature X. laevis females (Nasco, LM00535MX) were housed and maintained in the Stanford Aquatic Facility staffed by the Veterinary Service Center. To induce ovulation, frogs were primed 2–14 days before ovulation by the subcutaneous injection of 50U pregnant mare serum gonadotropin (PMSG; Sigma) at the dorsal lymph sac. Then, a second injection with 500 U human chorionic gonadotropin (hCG; Chorulon) induced frogs18h before ovulation. After the second injection, frogs were kept individually in 2ltrs of 1xMMR buffer at 17°C. Animal work was carried out in accordance with the guidelines of the Stanford University Administrative Panel on Laboratory Animal Care (APLAC).

Performed essentially as described^48^. Sperm DNA nuclei were added to CSF extract at a final concentration of 3×10^5^ sperm per 20 μL of extract and incubated in a water bath at 19°C for 60 min, flicking the tube every 15 min to prevent settling of the nuclei. After incubation, CSF extract was diluted with 1ml dilution buffer (1x BRB80 buffer (from 5x stock), 0.05% Triton X-100, 30% glycerol (from 100% stock), Add up add up to final volume with H2O, chill to 4°C before use) (5xBRB80: 400mM K-PIPES (from 0.5M stock at pH 6.8), 5mM MgCl2 (from 2M stock), 5mM K-EGTA (from 0.5 M stock), add up to final volume with H_2_O, filter and store at 4°C) and incubated for 5min on ice.

Then 1ml of fixation buffer (1x BRB80 buffer (from 5x stock), 0.05% Triton X-100, 30% glycerol (from 100% stock), 4% formaldehyde) was added and incubated for 5 min on ice. A sperm nuclei spin-down tube was prepared by layering custom-made^48^ platforms into a glass tube. 4ml of cushion buffer (1x BRB80 buffer (from 5x stock), 40% glycerol (from 100% stock), Add up to final volume with H_2_0, Chill to 4°C before use) was added to the tube and a poly-lysine coated coverslip was added to the platform. Sperm nuclei were carefully layered on top of the cushion buffer before being spun down at 3500 RPM for 20 min at 4°C. Buffers were aspirated and the coverslip was carefully removed from the platform. Coverslips were placed on parafilm and washed 3x with Abdil, before storing at 4°C in a humidified chamber until immunostaining.

Coverslips were incubated with primary antibodies (1:200 for Meikin and 1:500 for CENP-C antibodies) in Abdil for 1hr at RT, washed 3x with Abdil, then incubated with secondary antibodies (1:500) for 45 min at RT, washed 3x with Abdil and 2x with PBS. Coverslips were then mounted upside down in Vectashield with DAPI and sealed with nail polish. Slides were stored at −20°C until use.

### *X. laevis* micro-injection and meiotic timecourse

For micro-injection of stage VI X. laevis oocytes, mRNA to be injected (concentration ∼1μg/μl was spun down at max speed at 4°C for 30 min in a tabletop micro-centrifuge. Injection needles were pulled from glass capillaries (WPI) on a Sutter Needle puller (P-97). Needle tips were cut with a scalper to create an open tip. The needle was backloaded with mineral oil and attached onto a WPI nanolitre 2020 injector. mRNA was frontloaded into the needle. Oocytes were laid out onto custom made grooved trays (King’s Buildings workshop, University of Edinburgh) in 6 well culture plates (Corning) in 1xMBS media supplemented with 3% Ficoll 400 (Thermo Scientific) at room temperature. Oocytes were injected with 10ng of *xerp1C* mRNA or 2ng of *Meikin* (all constructs) mRNA. Oocytes were recovered in 1xMBS + 3% Ficoll 400 overnight at 18°C. For timecourses, oocytes were released from prophase arrest by transferring to 1xMBS + 3% Ficoll 400 with 10μg/ml final concentration of progesterone (Thermo Scientific) for 15 min at RT, then transferring again to 1xMBS + 3% Ficoll. GVBD was monitored by assessing the appearance of a white spot on the animal pole of the oocytes. For Meikin timecourses, samples were taken every 1 hr until 3.5 hrs, then every 30 min. For XErp1C timecourses, samples were taken every 1 hr. Samples were snap-frozen in liquid nitrogen and stored until use.

### Mice, ovaries collection, oocyte isolation and maturation

Sexually mature Female CD-1 IGS mice (Charles River, UK) were kept in the University of Edinburgh’s Bioscience and Veterinary Services (BVS) Ashworth facility under Home Office Licence PP3624594, Genetic and chromosome instability in the germline (licence holder: Ian Adams). Mice were euthanized by cervical dislocation under Schedule 1 of the Animals (Scientific Procedures) Act 1986 by licensed BVS staff. Ovaries were dissected by trained technicians and placed into pre-warmed (37^°^C) M2 handling medium (Sigma-Aldrich M7167), until transportation to the lab (5 minutes).

Oocytes were isolated by finely shredding ovaries and puncturing the follicles with a needle and forceps. Oocytes were kept in prophase-I arrest by addition of PDE3 inhibitor milrinone in M2 medium at a final concentration of 2μM at 37^°^C until experimental procedures. Based on the desired maturation stage, isolated oocytes were cleaned from the surrounded cumulus cells by repeated pipetting and then transferred to at least six drops of the pre-gassed maturation medium (M16) to washout the milrinone prior to incubation at 37^°^C and 5% CO_2_. Oocytes were monitored for meiosis resumption after milrinone washout by 1.5-2 hours by checking the Germinal vesicle breakdown (GVBD). For meiosis I and II, oocytes were kept in maturation medium after GVBD for 5 and 16 hours respectively.

### Mouse oocyte chromosome spreads and immunostaining

Zona-pellucida of meiosis I or II oocytes was removed by incubation in Acid Tyrode’s solution (Sigma-Aldrich) at 37^°^C and washed 3x in M2 media in 37^°^C. Oocytes were dropped in a 30µl drop of spread solution (1% formaldehyde (625μl of 16% stock in methanol), 3mM dithiothreitol (30 μl of 1M stock in H2O, make DTT fresh from powder every time), 0.15% Triton X-100 (75ul of 20% stock in H2O), Add up to 10ml H_2_O, pH to 9.2-9.3 with NaOH) on 12 well microscopy slides (Epredia) and allowed to lyse and spread. Spread slides were kept in an open humidified chamber overnight to air-dry. Dried spides were kept at −20^°^C until immunostaining

For immunostaining, slides were blocked for 10min with 30μl/well mammalian IF blocking buffer (3% bovine serum albumin (from powder), 0.05% Tween-20 (from 20% stock in H2O) in PBS, filter and store at 4^°^C), then primary antibodies were added 1:50 from stock in blocking buffer and incubated overnight at 4^°^C in a closed humidified chamber. Wells were washed 3x with 30μl/well of mammalian IF washing buffer (0.05% Tween (from 20% stock in H_2_O) in PBS) for 10 min at RT. 30µl of secondary antibodies 1:500 in blocking buffer was added to each well and incubated for 1hr at RT. Wells were washed once with 30µl mammalian IF wash buffer each for 5 min, then washed for 5 min in a bath of mammalian IF wash buffer. The outside edges of the slide was dried and 3 drops of Prolong antifade with hoechst were added before a coverslip was carefully laid over, allowing antifade solution to spread in all wells. Coverslips were sealed with nail polish then stored at 4^°^C until use.

### Trim away of HSPBAP1 in mouse oocytes and live imaging

Prophase-I arrested mouse oocytes were micro-injected using an Eppendorf Femtojet micro-injector in a milrinone containing M2-media at 37^°^C. mRNA encoding mouse SNAP-TRIM21was mixed 1:1 with a-HSPBAP1 antibody (Proteintech 1mg/ml) or rabbit IgG (1mg/ml) to a final needle concentration of mRNA of around 1μg/μl. Oocytes were live-imaged in milrinone free-M16 media supplemented with 500nM SiR-DNA and 500nM SPY555-tubulin at 37^°^C and 5% CO_2_.

For live-cell imaging: oocytes were mounted on a FluoroDish imaging dish, covered with oil and mounted on a Zeiss LSM 990 Airyscan microscope. Cells were imaged with a 647nm laser at 0.19% equivalent laser power and a 555nm laser at 4% laser power every 20 min with 2μm optical sections spanning the whole oocyte and imaging for at least 16 hours.

### Airyscan imaging of fixed mammalian samples

Fixed samples were imaged using a Zeiss LSM 980 or 990 Airyscan microscope equipped with an Airyscan 2 detector and a Plan-APO (63x/1.4 NA) oil objective (Zeiss UK, Cambridge) in Airyscan mode and ZEN pro software. 0.13-0.15 µM optical sections were used to image entire spindles or chromosome spreads. Samples were imaged with 405nm, 488nm, 561nm and 639nm lasers to detect Hoechst, alexa-488, alexa-555 and alexa-647 respectively.

### Donation of human oocytes to research and human oocyte culture

The NHS Research Ethics Committee approved this research project (Indicators of Oocyte and Embryo Development, 04/Q2802/26) and all work was conducted under a Research Licence from the Human Fertilisation and Embryology Authority (HFEA; R0155; Indicators of Oocyte and Embryo Development). Informed consent for donation of oocytes to research was provided by couples undergoing in vitro fertilisation (IVF) at the Edinburgh Fertility Centre and Department of Reproductive Endocrinology (EFC&DRE) at the Royal Infirmary of Edinburgh (NHS Lothian). Donations were optional and did not affect the treatment received. Couples were aware of the purpose of the research and were not provided compensation. All oocytes were considered unsuitable for treatment and would have otherwise been disposed of.

For collection of oocytes: Ovarian stimulation was induced using gonadotropins according to standard clinical protocols, either GnRH agonist or antagonist regimens. Oocytes were cultured and inseminated in the Vitrolife media suite at 37^°^C and 6% CO_2_. For couples undergoing ICSI treatment 6 hrs after collection, a maturation check was performed by a clinical embryologist. For couples undergoing IVF treatment: 17±1 hrs after insemination, a fertilisation check was performed by a clinical embryologist. Unmatured and unnfertilised oocytes were identified and collected from the clinic by licensed researchers 3-5 hours after the fertilisation check. Cells were transported in G-MOPS PLUS medium (Vitrolife) at 37^°^C in a portable incubator (K Systems) for approximately 15 min to the HFEA licensed research laboratory at the University of Edinburgh.

Oocytes were cultured in G-IVF PLUS (Vitrolife, #10136) under mineral oil (Merck, # 8042-47-5) at 37^°^C in 5% CO_2_, 6% O_2_ and 89% N_2_. Upon arrival in the research lab, the stage of the oocyte was confirmed by assessing the presence or absence of germinal vesicle and polar bodies.

### Immunofluorescence of human oocytes

Oocytes were fixated by first washing for 5 sec in PHEM buffer (60 mM PIPES, 25 mM HEPES, 10 mM EGTA, 4 mM MgSO4.7H2O, pH 6.9) with 0.25% Triton X-100 at 37^°^C and then fixated in PHEM buffer also containing 0.25% Triton X-100 and 4% formaldehyde for 30 minutes at room temperature. Cells were further permeabilised by first washing 3x in permeabilization buffer (PBS + 0.25% Triton X-100) followed by incubation in this buffer for 15 min. Cells were then washed 3x with wash buffer (PBS + 0.05% Tween-20) and stored in wash buffer for up to three weeks before immunostaining at 4dC.

For immunostaining, oocytes were first blocked with blocking solution (3% BSA and 0.05% Tween-20 in PBS) for 1hr at room temperature. Oocytes were then incubated with primary antibodies (a-HSPBAP1, a-UBR4 1:100 from stock, anti-CENP-C 1:200 from stock, anti-tubulin 1:500 from stock) in blocking solution overnight at 4^°^C. Primary antibodies were washed out in 3 washing steps with blocking buffer at room temperature with the final step lasting 1hr. Cells were transferred to secondary antibodies at room temperature (all 1:500 from stock in blocking solution), which were then washed out as above. Cells were immobilised in a resin drop of ProLong Gold hard-set antifade mountant with NucBlue (Hoechst) mixed 1:1 with PBS on a Fluorodish, which was allowed to harden overnight at room temperature in the dark. Cells were then stored at 4dC and imaged within two weeks.

### Trim away of HSPBAP1 in human oocytes and live imaging

Prophase-I stage human oocytes were micro-injected using an Eppendorf Femtojet micro-injector at 37^°^C. mRNA encoding human *TRIM-21eGFP* was mixed 1:1 with a-HSPBAP1 antibody (Genetex, 1mg/ml) or rabbit IgG (1mg/ml) to a final concentration of mRNA of around 1μg/μl. Oocytes were live-imaged in G-IVF media supplemented with 500nM SiR-DNA in at 37^°^C and 5% CO_2_.

For live-cell imaging: oocytes were mounted on a FluoroDish imaging dish, covered with oil and mounted on a Zeiss LSM 980 or 990 Airyscan microscope. Cells were imaged with a 647nm laser at 1% laser power to detect SiR-DNA every 15 or 20 min with 2μm optical sections spanning the whole oocyte for around 20-22 hours.

### Statistical information

Two-tailed t-test were performed with sample numbers indicated in figures or figure legends. Analysis of proteomics data was performed using the limma R-package^56^. Categorical data was compared with a Fisher’s exact test with sample numbers indicated in figures or figure legends. For all quantifications, medians are show with a horizontal bar. For boxplots, tails showing up the 95^th^ centile are shown.

---

**Extended Data Fig. 1.**
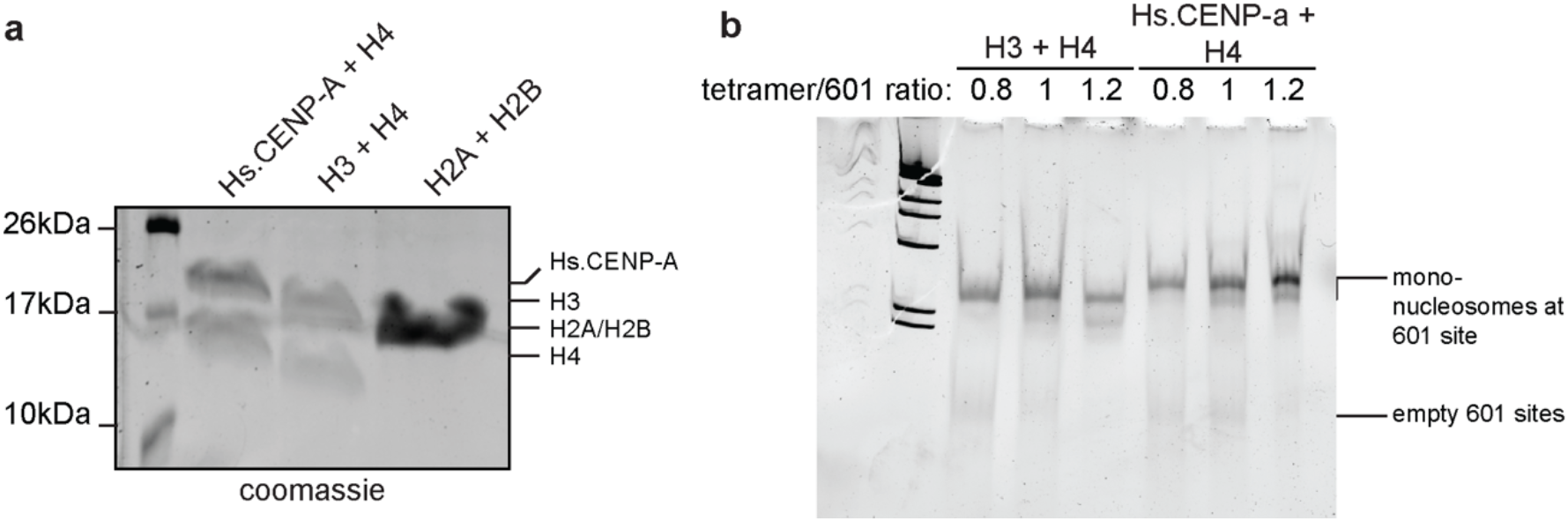
Synthetic chromatin preparation. **A)** Recombinant histones used for synthetic chromatin preparation. **B)** Example of synthetic chromatin preparation through dialysis. Different tetramer (CENP-A-H4 or H3-H4) to 601 site concentrations were used. For experiments CENP-A and H3 chromatin with similar ratios of nucleosome incorporation were used. For example tetramer/601 ratio of 1 in this experiment.

**Extended Data Fig. 2.**
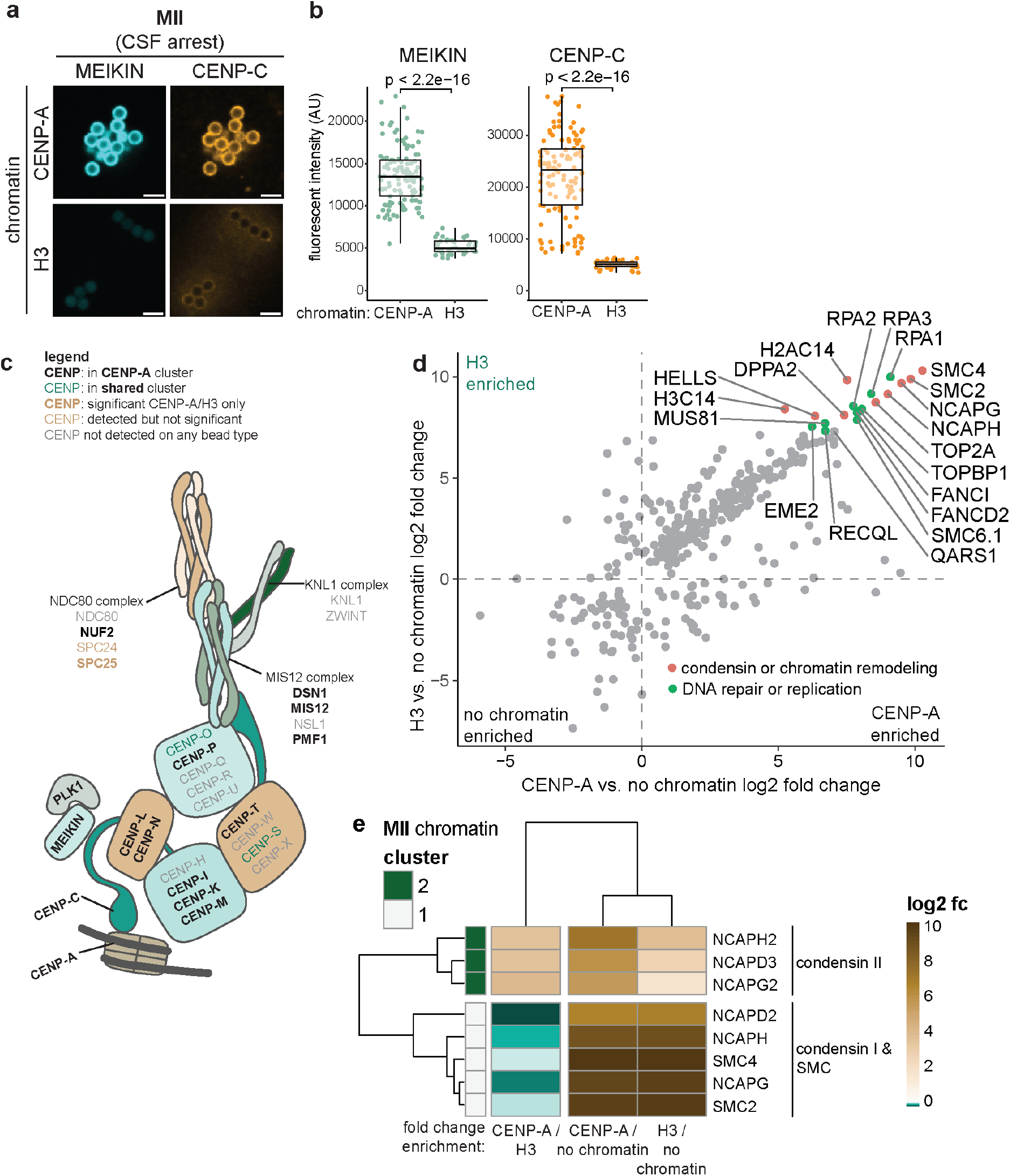
Meiosis II chromatin reconstitution. **A)** Widefield microscopy images of meiosis II synthetic chromatin beads immunostained for the indicated proteins. Scale bar represents 20μm. **B)** Quantification of experiment from data shown in (A). Each data point represents in individual bead. P-values calculated by t-test. **C)** Schematic overview of core kinetochore proteins detected on meiosis II centromeric chromatin. Different subcomplexes indicated through different colours. **D)** Plot showing the top 20 enriched proteins in the shared cluster of meiosis II reconstituted chromatin. X-axis represents the CENP-A/no chromatin enrichment, Y-axis represents the H3/no chromatin enrichment. Colour coded for manually annotated functions. **E)** Clustering of condensin subunits on meiosis II reconstituted chromatin based on enrichment levels.

**Extended Data Fig. 3.**
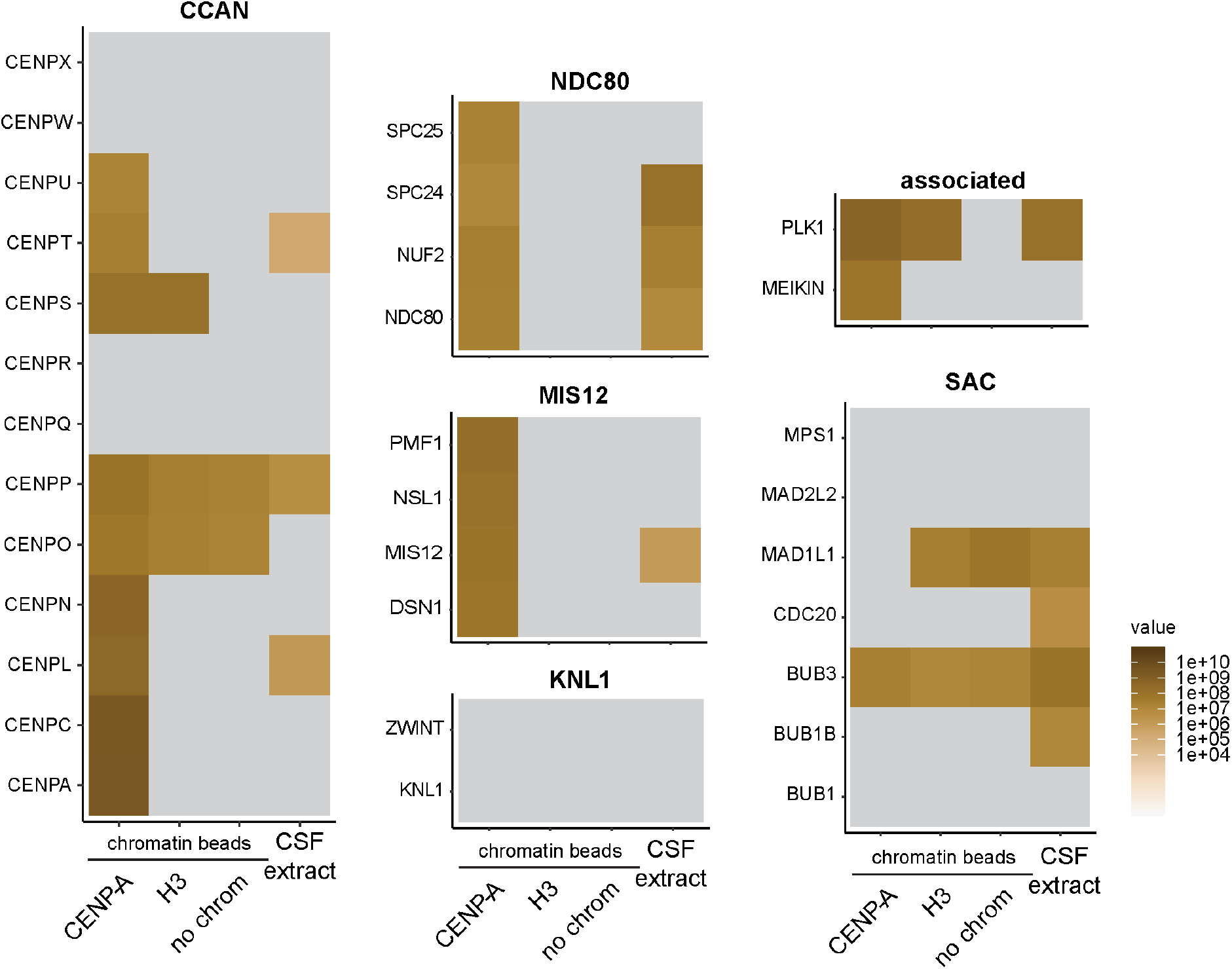
Detection of core kinetochore and kinetochore associated proteins on meiosis II reconstituted chromatin. Enrichment of proteins before imputation was averaged between repeats. Grey values indicate the protein was not detected in these samples. If a protein was detected in only a single repeat, the intensity value of this repeat is shown.

**Extended Data Fig 4.**
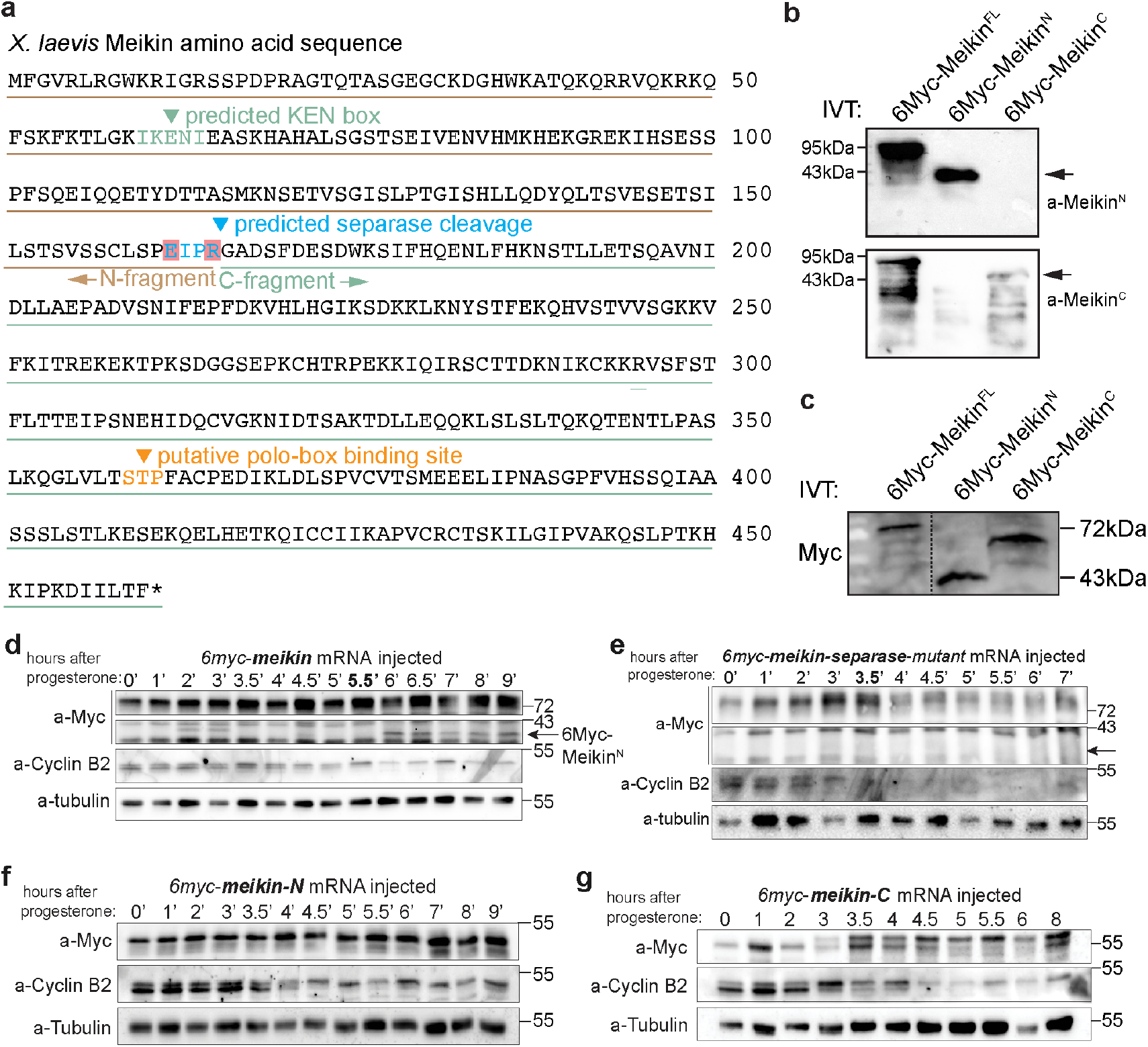
*X. laevis* MEIKIN can be proteolytically cleaved. **A)** Amino-acid sequence of *X. laevis* MEIKIN. With indicated sites of interest. The KEN-box was predicted using the eukaryotic linear motif resource (http://elm.eu.org/). The potential separase cleavage site and STP site were annotated based on alignment with the mouse sequence based on the separase cleavage site and STP sites identified in Maier et al. (2021). Residues highlight in red are mutant in the *X. laevis* MEIKIN-separase-mutant. **B)** Detection by western blot of in vitro translated (IVT) *X. laevis* MEIKIN constructs with specific MEIKIN^N^ and MEIKIN^C^ antibodies. Arrows indicate the location of the respective MEIKIN^N^ and MEIKIN^C^ fragments. **C)** Detection by western blot of in vitro translated *X. laevis* MEIKIN constructs with a-Myc antibodies. **D)** Example of a western blot time course of 6myc-MEIKIN mRNA injected *X. laevis* oocytes. Membrane was stripped for each different incubation. N=5 **E)** Example of a western blot time course of 6myc-MEIKIN-separase-mutant mRNA injected *X. laevis* oocytes. Membrane was stripped for each different incubation. N=3 **F)** Example of a western blot time course of 6myc-MEIKINN mRNA injected *X. laevis* oocytes. Membrane was stripped for each different incubation. N=4 **G)** Example of a western blot time course of 6myc-MEIKINC mRNA injected *X. laevis* oocytes. Membrane was stripped for each different incubation. N=3

**Extended Data Fig. 5.**
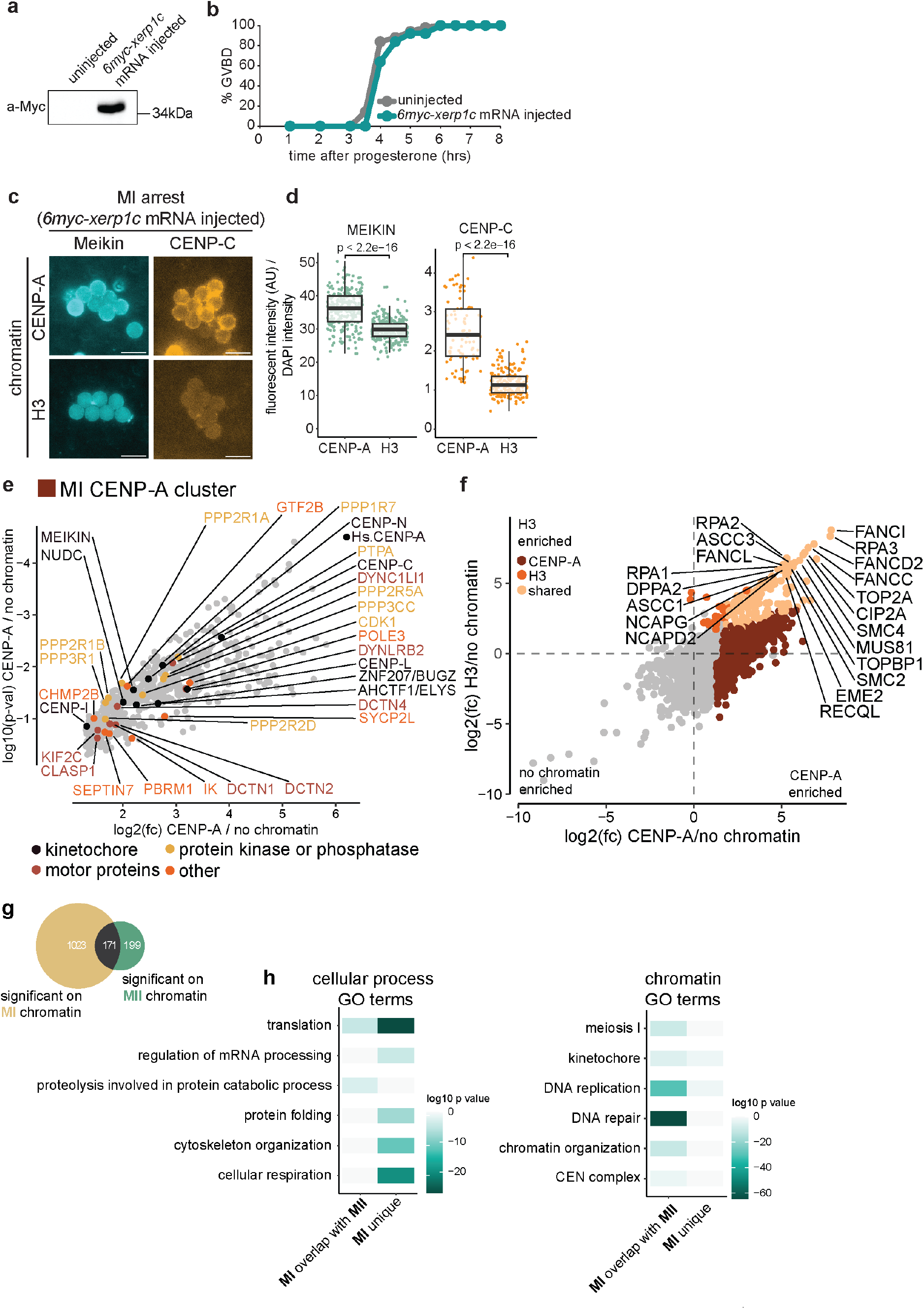
Meiosis I reconstituted chromatin. **A)** Example of expression of 6Myc-Xerp1C in prophase arrested oocytes at the start of a time course. **B)** Example of monitoring of GVBD rate of uninjected and *6myc-xerp1c* mRNA injected oocytes through the timecourse **C)** Immunofluorescence images of meiosis I reconstituted chromatin on beads stained for indicated proteins. Scale bar represents 20uM **D)** Quantification of data shown in (C). Each data point represents in individual bead. P-values calculated by t-test. **E)** Plot showing all the proteins in the meiosis I CENP-A cluster. Y-axis represents CENP-A/no chromatin fold change and the X-axis the p-values. Centromere and kinetochore associated proteins were labelled and colour coded for protein function. **F)** Plot showing the different protein clusters same in Fig. 2C, but with the top 20 proteins in the shared cluster labelled. **G)** Venn diagram showing the overlap between all proteins that were significant on any type of bead on meiosis I and meiosis II reconstituted chromatin. **H)** GO-term analysis of the proteins that are unique on MI chromatin (n=1023) and that overlap with MII chromatin (n=171).

**Extended Data Fig. 6.**
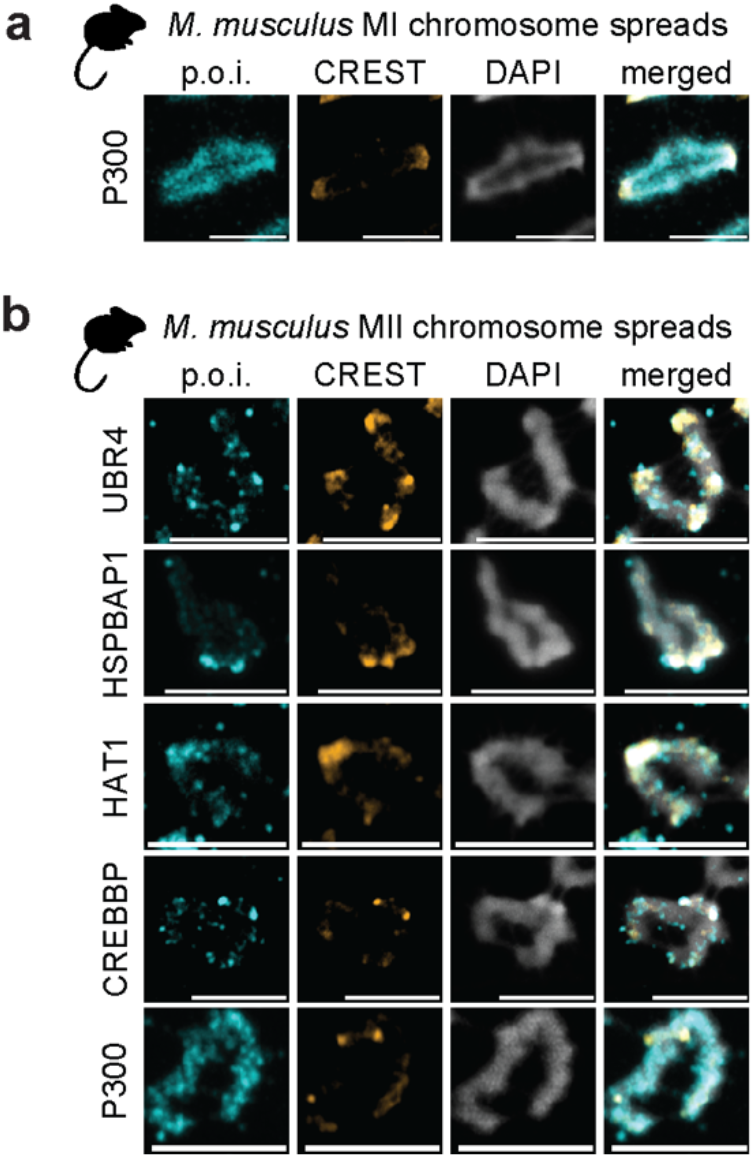
Conservation of novel kinetochore proteins to mammalian meiotic centromeres. **A)** Representative mouse meiosis I chromosome spreads immunostained with indicated antibodies. Scale bar represents 5μm. **B)** Representative mouse meiosis II chromosome spreads immunostained with indicated antibodies. Scale bar represents 5μm.

**Extended Data Fig. 7.**
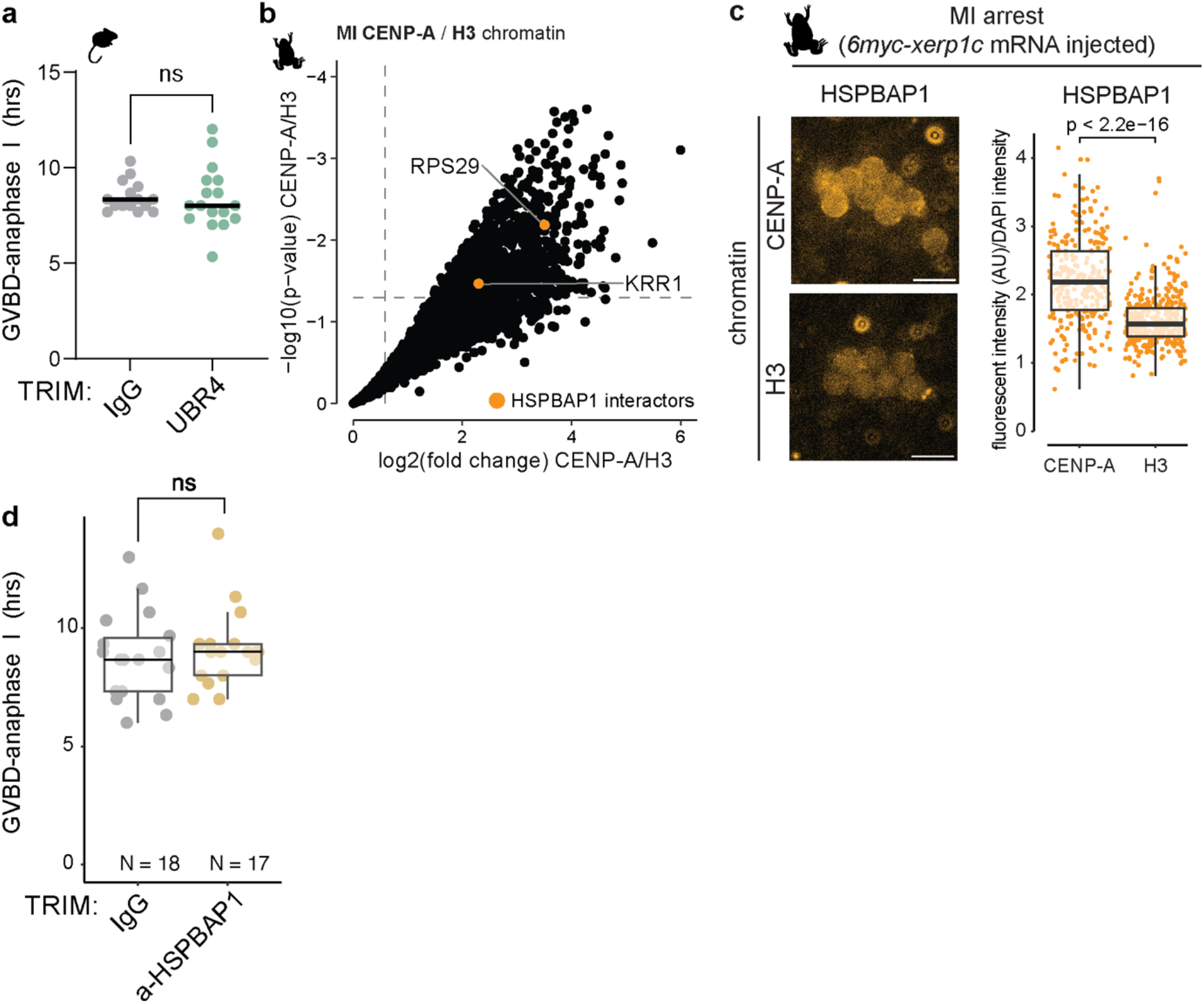
UBR4 and HSPBAP1 function in mouse meiosis. **A)** Timing of meiosis I duration in mouse oocytes injected with *Mm.TRIM-SNAP* mRNA and either rabbit IgG or a-UBR4 antibody (N=17 for IgG and UBR4 TRIM from 3 experiments). P-value calculated by t-test. **B)** Volcano plot showing CENP-A enrichment of proteins on meiosis I reconstituted chromatin. Colour coded are HSPBAP1 interactors detected on CENP-A chromatin. Grey dashed lines represent a p-value of 0.05 and a fold change of 2. **C)** Chromatin reconstitution in meiosis I arrested (*6myc-xerp1c* mRNA injected) oocytes extract. Left panel: immunofluorescence images of chromatin beads stained with a-HSPBAP1 antibody. Right panel: HSPBAP1/DAPI fluorescent intensity. Each dot represents one bead. P-value calculated by t-test. **D)** Timing of meiosis I duration in mouse oocytes injected with *Mm.TRIM-SNAP* mRNA and either rabbit IgG or a-HSPBAP1 antibody. P-value calculated by t-test.

