## Supplemental Table 1 for "Meiotic kinetochore reconstitution identifies chromosome segregation factors"

### **Supplementary Table 1**

### Reagents

| **Reagent** | **Supplier** | **Catalog Number** |
| --- | --- | --- |
| Biotin-14-dATP | Life Technologies Ltd | 19524016 |
| Biotin-16-dUTP | Jena Bioscience | NU-803-BIO16-S |
| dCTP | Invitrogen | 10217016 |
| dGTP | Thermo Fisher | 10218014 |
| Klenow Fragment (3’-5’ Exo-) | NEB | M0212M |
| Streptavidin-FITC | Merck | S3762-.1MG |
| Dynabeads M-270 streptavidin | Invitrogen | 653-05 |
| Collagenase B | Merck | 11088831001 |
| Cytochalasin B | Scientific Laboratory Supplies Ltd | C6762-5MG |
| Ficoll 400 | Fisher Scientific | 11469597 |
| Cytochalasin D | Stratech Scientific | B6645-APE-1mg |
| rapigest | Fisher Scientific | 10103214 |
| poly-lysine | Merck | P1524-25MG |
| Vectashield with DAPI | Vector | H1200 |
| PMSG | Intervet Chorulon |  |
| HCG | Intervet Chorulon |  |
| ProLong Gold with NucBlue | Thermo Fisher Scientific | P36931 |
| M2 media | Sigma-Aldrich | M7167 |
| M16 media | Sigma-Aldrich | M7292 |
| milrinone | Sigma-Aldrich | M4659 |
| Acid Tyrode’s solution | Sigma-Aldrich | T1788 |
| Bovine serum albumin | Sigma-Aldrich | A9647-100G |
| Rabbit IgG | Sigma-Aldrich | #12-370 |
| SiR-DNA | Tebubio | SC007 |
| SPY555-tubulin | Tebubio | SC203 |
| G-MOPS media | Vitrolife | #10130 |
| G-IVF media | Vitrolife | #10136 |
| Mineral oil | Sigma | M8410-1L |
| Ficoll 400 | Thermo Scientific | B22095.18 |
| Progesterone | Thermo Scientific | P8783-1G |

### Antibodies

| **Target** | **Host** | **Reactivity** | **Source** |
| --- | --- | --- | --- |
| a-CENP-C | Rabbit | *X. laevis* | Ana Losada |
| a-Meikin-C | Sheep | *X. laevis* | This work |
| a-Meikin-N | Sheep | *X. laevis* | This work |
| a-Cyclin B2 | sheep | *X. laevis* | EXRC |
| a-Myc | mouse | Human | antibodies.com A278887 |
| a-HSPBAP1 | rabbit | Human, mouse | Genetex GTX116828 |
| a-HSPBAP1 | rabbit | Human, mouse | Proteintech 27631-1-AP |
| a-UBR4 | rabbit | Human, mouse | Invitrogen 15722251 |
| a-CREBBP | rabbit | Mouse | Proteintech 22277-1-AP |
| a-HAT1/KAT1 | rabbit | Mouse | Abcam AB194296 |
| a-P300/KAT3B | rabbit |  | Abcam AB275378 |
| a-NPM2 | rabbit |  | Abcam AB243544 |
| a-RBBP7 | rabbit |  | Abcam AB259957 |
| a-CENP-C | Guinea pig | Human | MBL PD030 (discontinued) |
| a-𝛂-tubulin | mouse | Human, mouse | Sigma T6074 |
| a-GFP | rabbit | *Aequorea victoria* | Abcam ab290 |
| a-rabbit a488 | goat | rabbit | ThermoFisher Scientific A-11008 |
| a-mouse a555 | goat | mouse | Thermo Fisher Scientific A-21422 |
| a-human a555 | goat | human | A48277 |
| a-guinea pig a647 | goat | guinea pig | Thermo Fisher Scientific A-21450 |

### Materials

| **Item** | **Supplier** | **Catalogue number** |
| --- | --- | --- |
| Dounce homogeniser |  |  |
| HiTrap Q FF column 5ml | Cytiva | 17515601 |
| HiTrap S FF column 5ml | Cytiva | 17515701 |
| CHTII Ceramic Hydroxyapatite 60ml | Bio-Rad | #1572000 |
| HiTrap SP FF column 1ml | Cytiva | 17505401 |
| Superdex 200 16/60 |  |  |
| Ultra clear centrifuge tubes | Beckman Coulter | 344090 |
| SW55-Ti Adaptors for ultra clear tubes | Beckman Coulter | 356860 |
| fluorodish | WPI | FD35-100 |
| 12 well microscopy slides | Epredia | X1XER302W |
| Femtojet micro-injector | Eppendorf |  |
| Capillaries | WPI | 504949 |
| Nanoliter 202 injector | WPI |  |
| 6-well culture plates | Corning | 3516 |

### Plasmids

| **ID** | **Backbone** | **Use** | **Species** | **Source** |
| --- | --- | --- | --- | --- |
| ASP341 | pET3aTr | Xl.H2A | X. laevis | Ref^1^ |
| ASP342 | pET3aTr | Xl.H2B | X. laevis | Ref^1^ |
| ASP343 | pET3aTr | Xl.H3 | X. laevis | Ref^1^ |
| ASP1123 | pET3aTr | Xl.H4 | X. laevis | Ref^1^ |
| ASP2773 | pST39 | Hs.CENP-A, myc-Xl.H4 coexpression | X. laevis/ H. sapiens | Ref^2^ |
| ASP696 | pUC18 | 18x 601 | - | Ref^3^ |
|  | pCS2 | 6Myc-xErp1-491-651, T545A T551A | X. laevis | Thomas Mayer |
| AMp2009 | pDEST17 | 6His-Xl.Meikin-N | X. laevis | This work |
| AMp2010 | pDEST17 | 6His-Xl.Meikin-C | X. laevis | This work |
| AMp2350 | pGEMHE | 6Myc-Xl.Meikin-FL | X. laevis | This work |
| AMp2351 | pGEMHE | 6Myc-Xl.Meikin-C | X. laevis | This work |
| AMp2392 | pGEMHE | 6Myc-Xl.Meikin-N | X. laevis | This work |
| AMp2560 | pGEMHE | 6Myc-Xl.Meikin-separase-mut | X. laevis | This work |
| Amp2582 | pCDNA3 | eGFP-TRIM21 | human | This work |
|  | pGEM | SNAP-TRIM21 | mouse | Binyam Mogessie Addgene #196181 |
